# Integrated physiological markers of drought tolerance and yield stability in cotton under deficit irrigation

**DOI:** 10.64898/2026.05.31.729145

**Authors:** Noemi Tortorici, Xuejun Dong, Thiago F. Duarte, Nicolò Iacuzzi, Uzair Ahmad, Teresa Tuttolomondo

## Abstract

The selection of drought-resistant cotton genotypes with high productivity and improved water-use efficiency is an increasingly pressing challenge in arid and semi-arid cotton-growing regions, where climate variability is intensifying water scarcity. This study evaluated two widely cultivated cotton varieties in Texas, NG 4190 B3XF and ST 4990 B3XF, under full and moderate deficit irrigation to identify physiological markers associated with contrasting yield responses under water limitation.

A comprehensive set of physiological traits was assessed, including plant water status, leaf gas exchange, carbon isotope composition (δ13C), total nitrogen and C/N ratio, chlorophyll fluorescence, light and CO□ response curves, and canopy temperature. While no yield differences were observed under full irrigation, moderate water deficit resulted in stable seed and fiber yield in NG 4190, but significant yield reduction in ST 4990.

Yield differences were not explained by instantaneous gas exchange or direct biochemical limitations of photosynthesis, but rather by integrated physiological behavior over time. Key discriminating traits included midday relative water content (RWC), photosynthetic light-response parameters (α and P_m_), stomatal optimization parameter (g□), and δ13C. NG 4190 exhibited higher RWC, more negative δ13C, and higher g□ values, indicating a less conservative stomatal regulation strategy that supported sustained carbon assimilation under water stress.

These findings provide insight into the physiological mechanisms underlying cotton performance under drought and support the use of integrated physiological markers for the selection of resilient genotypes in water-limited environments.

## 1. Introduction

The selection of high-yielding cotton genotypes represents an increasingly challenging task, despite significant progress in breeding programs (Passioura et al. 2006; Ghorbanzadeh et al. 2025; Yang et al. 2023;), particularly in the context of growing water scarcity and stringent environmental sustainability constraints (Texas Water Development Board 2022). Cotton (*Gossypium hirsutum* L.) is a crop of major economic importance in the United States, with a production of approximately 3 million t; among producing states, Texas ranks first, contributing approximately 30% of national production (USDA-FAS 2026). In this context, ensuring long-term yield stability and preserving Texas’s role as the primary national production hub, while reducing pressure on freshwater resources, requires strategies to improve water use efficiency.

Addressing this challenge requires, among other strategies, direct intervention at the plant level. Key plant-based approaches include: i) the study of natural genetic variability and the relationships between traits influencing water use efficiency and yield under water-limited conditions; and ii) the transgenic manipulation of genes involved in the regulation of photosynthetic apparatus, stomatal conductance (g_s_), mesophyll conductance (g□), and plant hydraulics (Leakey et al. 2019). In both cases, the ultimate goal is to optimize photosynthetic processes to sustain high carbon assimilation even under limited water availability. However, photosynthetic efficiency results from a complex interplay of physiological, biochemical, and hydraulic mechanisms, whose integration and regulation under water stress are not yet fully understood (Baker et al. 2006; Passioura et al. 2006; Lamour et al. 2022).

Given this complexity, reliably predicting drought tolerance remains difficult. Although measuring a single trait is less time-consuming, multiple traits are often necessary to explain and predict crop drought tolerance (Ko and Piccinni 2009; Templer et al. 2017; Li et al. 2022).

Integrative indicators that combine plant water status and carbon assimilation over time are more effective for distinguishing genotypic responses (de Brito et al. 2011; Sinclair 2012; Medrano et al. 2015). Moreover, commonly used instantaneous physiological measurements, such as instantaneous leaf gas exchange, may not be sufficient to explain differences in productivity between genotypes, particularly under moderate water stress, highlighting the need for indicators that capture plant physiological responses during the growing season (Medrano et al. 2015; McAusland et al. 2016).

Major drought tolerance mechanisms associated with high yield and productivity stability include traits such as osmotic adjustment, antioxidant metabolism, membrane stability, leaf morphological characteristics (e.g., increased leaf thickness), the ability to maintain tissue water potential, and high stomatal conductance (Niinemets et al. 2001; Templer et al. 2017; Li et al. 2022). Among these traits, stomatal conductance plays a central role in regulating gas exchange. Stomata have been estimated to limit photosynthetic assimilation by up to 20%, with potentially significant effects on productivity (McAusland et al. 2016). Although high stomatal conductance may reduce intrinsic water use efficiency (iWUE) due to increased transpiration (Jones 2007), it simultaneously promotes greater CO□ flux to the carboxylation sites, supporting photosynthesis and yield even under under water-limited conditions (de Brito et al. 2011; Luo et al. 2016). This trade-off between carbon assimilation and water use can be partially mitigated through more efficient regulation of stomatal, mesophyll, and hydraulic traits (Jones 2007; Medlyn et al. 2011; Buckley and Farquhar 2017; Sunoj et al. 2025).

The present study aimed to identify the cotton genotype more tolerant to moderate water stress among two widely cultivated varieties in Texas (NG 4190 B3XF and ST 4990 B3XF), which exhibit contrasting productivity responses. In addition, the study investigated the physiological mechanisms underlying yield reductions under water deficit, and identified physiological traits that can serve as reliable markers for field-based selection of drought-tolerant varieties. Our hypothesis was that, under moderate water stress, differences in yield stability between genotypes were primarily associated with variation in leaf photosynthetic traits and stomatal behavior.

To test this hypothesis, a range of physiological traits related to plant water status, gas exchange, and photosynthetic performance were analyzed, including carbon isotope composition (δ^13^C), total N and C/N ratio, chlorophyll fluorescence, light and CO_2_ response curves, and canopy temperature, to characterize genotypic differences.

The results provide insights into the physiological responses of cotton to water stress and indicate that some of these traits may serve as reliable markers for selecting high-performing genotypes, thereby supporting breeding strategies aimed at maintaining yield stability and improving water use efficiency in semi-arid environments.

## 2. Materials and Methods

### 2.1. Experimental site, crop management and experimental design

The study was conducted under open-field conditions at the Texas A&M AgriLife Research and Extension Center in Uvalde, Texas, USA (29°13′03″ N, 99°45′26″ W; 283 m a.s.l.), from April to September 2025. The site is characterized by a semi-arid climate and a clay soil (fine-silty, mixed, active, hyperthermic Aridic Calciustoll) of the Uvalde series (USDA 1970). Figure 1 summarizes meteorological data, including temperature, precipitation, and vapor pressure deficit (VPD), collected throughout the growing season using an on-site weather station.

**Figure 1.**
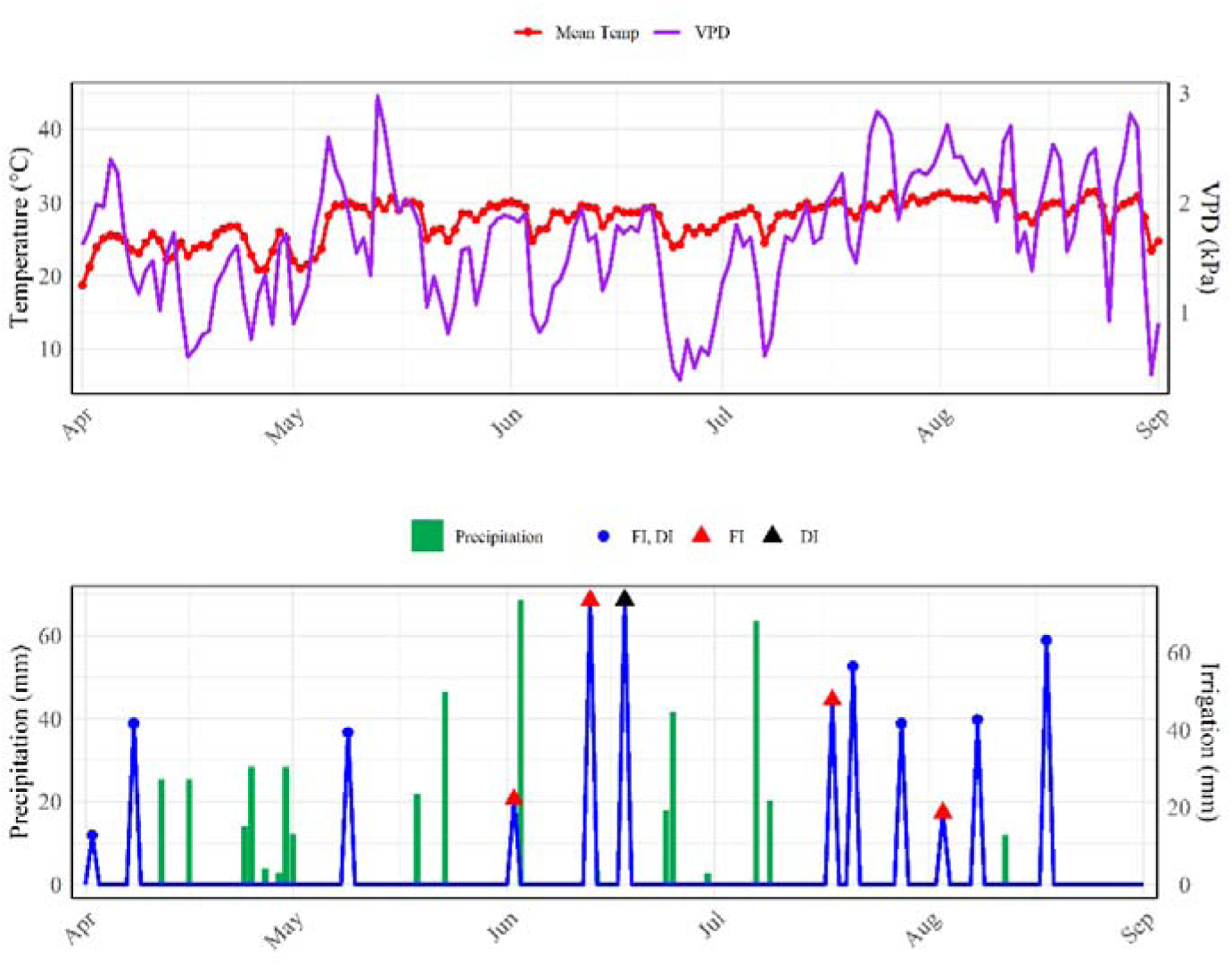
Average daily temperature (Mean Temp, °C), precipitation (mm), and vapor pressure deficit (VPD, kPa) measured by an on-site weather station during the 2025 growing season at Uvalde, Texas. Also shown are the amounts of irrigation (mm) applied in individual events through a drip system. FI = full irrigation; DI = deficit irrigation. Labels “FI, DI” denote irrigation events applied to both treatments, while “FI” and “DI” denote events applied only to the respective treatment.

Two cotton (*Gossypium hirsutum* L.) commercial cultivars, NG 4190 B3XF and ST 4990 B3XF, were selected based on their contrasting responses to water stress observed in a previous field survey study conducted in the same experimental environment (X. Dong, unpublished).

The experimental field was level and had been cultivated with a mixture of cover crops during the previous growing season. Mineral fertilization was applied to ensure adequate nutrient availability, consisting of 100 kg N ha^-1^ and 26 kg P□O□ ha^-1^. No potassium fertilizer was applied because soil K levels were considered sufficient. Approximately three weeks before planting, the field was irrigated with 50 mm of water using a linear sprinkler irrigation system.

The cotton field was planted on 8 April 2025 using a six-row vacuum planter. After crop emergence, irrigation was supplied through surface drip irrigation tapes, with one drip line installed per crop row. Crop management followed the recommended practice for the region. Harvest was performed manually on 3-4 September 2025, when approximately 90% of bolls had reached maturity (BBCH 89).

The experimental protocol evaluated two factors: (i) variety, with two levels (NG 4190 B3XF; ST 4990 B3XF), and (ii) irrigation regime, with two levels (FI: full irrigation, soil water content restored to 100% of field capacity; DI: deficit irrigation, soil water content restored to 75% of FI).

A split plot design was used, with irrigation as the main plot and genotype as the subplot, and three replicates per treatment. The total experimental area (∼0.405 ha) was divided into twelve plots. Each plot consisted of four rows, 60 m in length, with a row spacing of 0.76 m.

### 2.2. Irrigation management

Soil water content (SWC) was monitored in all plots from 0 to 100 cm soil depth at intervals of 10, 20, 40, 60, 80 and 100 cm, using a CPN 503 Elite Hydroprobe (InstroTek, Inc., Research Triangle Park, NC, USA). Measurements were taken through aluminum access tubes installed at the center of each experimental plot. The neutron probe was calibrated using the two-point method as recommended by the manufacturer. SWC was measured at approximately 14 day intervals. Soil water storage was expressed as equivalent water depth (mm).

SWC at field capacity (θ_fc_) and permanent wilting point (θ_wp_) were 0.328 and 0.179 m^3^ m^-3^, respectively, as previously determined using laboratory pressure plate methods. For irrigation scheduling, total available water (TAW, mm) in the 0–40 cm soil layer was calculated as:

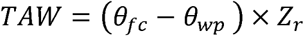

where Z_r_ is the effective rooting depth (400 mm). For the full irrigation treatment, the allowable depletion threshold (D_p_, mm) was calculated following Allen et al. (1998):

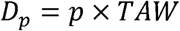

where *p* is the depletion fraction, set to 0.5 according to Allen et al. (1998). Irrigation was applied when soil water deplation in the 0–40 cm soil layer approached D_p_, in order to restore soil moisture to field capacity. The deficit irrigation treatment received approximately 75% of the irrigation volume applied to the full irrigation treatment. Total seasonal irrigation amounts for the FI and DI treatments are reported in Table 1.

**Table 1.** Total amounts of irrigation and rainfall received by the cotton crop during the growing season, from 8 April (sowing) to 4 September 2025 (harvest).

| Treatment | Irrigation<br>(mm) | Rain<br>(mm) | Irrigation + rain<br>(mm) |
| --- | --- | --- | --- |
| FI | 460 | 438 | 898 |
| DI | 342 | 438 | 780 |

Note: FI = full irrigation; DI: deficit irrigation.

### 2.3. Crop measurements

#### 2.3.1. Leaf water potential, relative water content, and leaf area index

Leaf water potential (LWP, MPa) and relative water content (RWC, %) were measured at predawn and midday on five clear days (D1 = 9 June; D2 = 25 June; D3 = 10 July; D4 = 1 August; D5 = 28 August). For each measurement, three healthy leaves from the fourth node of randomly selected plants in each of the 12 plots were sampled.

LWP was determined using a pressure chamber (Model 615, PMS Instrument Company, Albany, OR, USA), with the high pressure provided by compressed pure nitrogen gas. After LWP measurement, each leaf was immediately placed in a sealable plastic bag of known weight, kept on ice in the dark, and promptly transported to the laboratory for RWC determination, following the method of Barrs and Weatherley (1962). Fresh weight (FW) was recorded, then leaves were floated in distilled water for 24 h at cool temperature, gently blotted, and weighed to obtain turgid weight (TW). Finally, leaves were oven-dried at 70 °C until constant weight to determine dry weight (DW).

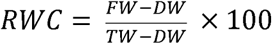

Leaf area index (LAI) was measured in all 12 plots once every two weeks throughout the growing season using a LI-2200C Canopy Analyzer (LI-COR Biosciences, Lincoln, NE, USA). In each plot, the canopy was scanned five times along a 15-m transect near the center of the plot, and the mean of the five measurements was recorded as the plot LAI.

#### 2.3.2. Gas exchange measurements

Diurnal leaf gas exchange was measured on the same five clear days (D1–D5) using a LI-6400XT Portable Photosynthesis System (LI-COR Biosciences, Lincoln, NE, USA). Measurements were performed 3-4 times per day between 08:00 and 14:30 h on three fully sun-exposed leaves from the fourth node of a randomly selected representative plant in each plot. A 2 × 3 cm leaf chamber equipped with a 6400-02B LED light source was used. Chamber conditions were set as follows: photosynthetic photon flux density (PPFD) was adjusted to ambient light conditions using measurements from an external quantum sensor (LI-190SA) at the beginning of each measurement period; reference CO□ concentration was set to 450 µmol mol^-1^; leaf temperature and relative humidity were not actively controlled; flow rate was set to 500 µmol s^-1^. Water vapor was partially scrubbed using fresh desiccant, typically resulting in a chamber relative humidity of ∼50-60% when the cuvette was fully covered by the leaf.

Gas exchange data from each genotype were used to separately fit the Medlyn et al. (2011) stomatal conductance (g_s_) model:

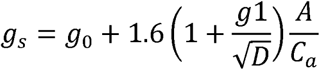

where g_s_ is stomatal conductance (mol m^-2^ s^-1^), g□ is a fitted parameter (kPa^0.5^), A is the net CO□ assimilation rate (µmol m^-2^ s^-1^), D is vapor pressure deficit (kPa), and C□ is the atmospheric CO□ concentration at the leaf surface (µmol mol^-1^). The intercept parameter g_0_ was fixed to zero. The parameter g□ was estimated by fitting observed g_s_ to model-predicted values using nonlinear regression in Minitab Statistical Software 22 (Minitab, LLC, State College, PA, USA).

#### 2.3.3. Photosynthetic capacity and CO□-based light response

Photosynthetic capacity and light response were assessed by measuring A/C□ curves and CO□-based rapid light response curves (RLC), respectively, on fully expanded leaves from the fourth node.

A/C□ curves were measured on 12 leaves (three per plot from four plots representing two cultivars under two irrigation treatments) on 9 and 13 August, between 10:00 and 14:00 h, following the recommended protocol of LI-6400XT Portable Photosynthesis System. Chamber conditions were set as follows: PPFD was maintained at 1500 μmol m^-2^ s^-1^; flow rate was set to 500 μmol s^-1^; leaf temperature was held constant at 30 °C, corresponding to the temperature observed at 10:00 h; and water vapor mole fraction was adjusted to maintain the chamber relative humidity at approximately 50-60%. During the A/C_i_ curve measurement, chamber CO_2_ concentration was adjusted to change in the sequence of the following steps: 400, 300, 200, 100, 50, 440, 440, 600, 800 µmol mol^-1^.

CO□-based RLC were measured on 9 August, between 09:00 and 13:00 h, on the same plots, following the LI-COR protocol. During RLC measurement, PPFD varied from 1500 to 0 μmol m^-2^ s^-1^ in nine steps: 1500, 800, 500, 300, 200, 100, 50, 20, 0 μmol m^-2^ s^-1^, while the reference CO_2_ concentration was mantained at 450 µmol mol^-1^. All other chamber conditions were kept at the values used for the A/C□ curve measurements.

Parameters of the A/C□ curves were estimated using the online fitting tool Leaf Web (Gu et al. 2010), which fits the Farquhar–von Caemmerer–Berry biochemical model of photosynthesis (Farquhar et al. 1980).

Light response curves were fitted using a non-rectangular hyperbola model (Thornley 1998):

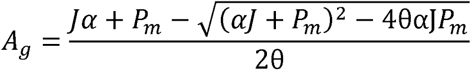

where A_g_ is gross photosynthetic rate (μmol CO□ m^-2^ s^-1^), J is PPFD (μmol photons m^-2^ s^-1^), α is apparent quantum yield, P_m_ is maximum gross photosynthetic rate, and *θ* is the photosynthesis sharpness parameter of the light response curve. The suitability of this model for describing photosynthesis-light response curves has been discussed previously (Thornley 2002). Gross photosynthetic rate was calculated by adding dark respiration (measured as net CO□ exchange rate in darkness, expressed as an absolute value) to the net photosynthetic rate measured at different light levels prior to curve fitting.

#### 2.3.4. Chlorophyll fluorescence

The maximum photochemical efficiency of photosystem II (PSII; F_v_/F_m_) was measured on August 1 and 28, between 09:00 and 09:30 h on three fully expanded leaves per plot (fourth node). Leaves were dark-adapted for 40 min using the instrument clips, and measured with an OS1p fluorometer (Opti-Sciences Inc., Hudson, NH, USA). After dark adaptation, the fiber-optic probe was positioned on the leaf surface, and the saturating pulse protocol was applied. Measuring light intensity was set to 10% of the maximum (≈0.1-1 μmol m^-2^ s^-1^) to determine minimal fluorescence (F□), and the saturating pulse was set at 75% of maximum output (≈8250 μmol m^-2^ s^-1^, 0.8 s duration) to obtain maximal fluorescence (F□). F_v_/F_m_ was calculated as:

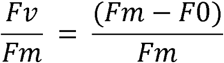

Fluorescence-based RLC were measured on 9 and 28 August, between 11:00 and 11:30 h, on 12 fully expanded leaves (three per plot) from the same plots used for A/C_i_ and CO_2_-based rapid light response measurements. A PAR clip was used, and measurements followed OS1p manufacturer guidelines. The RLC consisted of 10 increasing actinic light steps (10 s each), with saturating pulses (100%, 1 s) applied at each step. Key parameters of the RLC were estimated using the Jassby and Platt (1980) model:

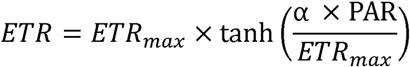

where α is the initial slope (α_PSII_, photosynthetic quantum efficiency), ETR_max_ is the maximum electron transport rate, and I_k_ = ETR_max_/ α, is the light saturation parameter.

#### 2.3.5. Canopy temperature

Canopy temperature (T_c_) was measured on the same five clear days (D1–D5) as RWC, LWP, and gas exchange measurements. Measurements were conducted three times per day between 08:00 and 14:00 h. For each measurement, three healthy plants were randomly selected in each of the 12 plots. T_c_ was recorded using a hand-held infrared radiometer (MI-210, Apogee Instruments Inc, UT, USA). For each replicate, canopy temperatures were measured from four different angles, spaced 90° apart, and subsequently averaged. The radiometer was positioned to minimize soil interference and to maximize canopy coverage within the field of view. ΔT was calculated as the difference between canopy temperature (Tc) and air temperature (Tair) measured at the same time of canopy temperature acquisition by the on-site weather station, for each measurement date and time.

#### 2.3.6. Leaf carbon isotope composition, total N, and C/N ratio

Leaves collected for midday RWC measurements on each of the five sampling dates were subsequently used for carbon isotope ratio (δ^13^C), total nitrogen (TKN), and C/N ratio analyses. Carbon isotope composition and total carbon content were determined at the UC Davis Stable Isotope Facility, Department of Plant Sciences, Davis, CA, USA. Dried leaves were finely ground with a mortar and pestle to obtain a homogeneous sample. For each date and each plot, 2-3 mg of dry material was weighed and analyzed for δ^13^C and total carbon content. Samples were combusted in an elemental analyzer (EA-IRMS) system to convert organic carbon to CO□. Residual gases were removed, and CO□ was separated and introduced into an isotope ratio mass spectrometer for δ^13^C measurement, following the standard methodology of the UC Davis Stable Isotope Facility. All δ^13^C measurements were calibrated against the international reference material VPDB and quality-controlled using in-house reference materials to ensure accuracy and precision.

Total nitrogen (TKN) was determined separately using the automated discrete colorimetry method (O’Dell 1993) at the Texas A&M AgriLife Research and Extension Center in Uvalde, Texas, USA. The C/N ratio was calculated from total carbon and total nitrogen concentrations.

#### 2.3.7. Yield and fiber quality

Seed cotton yield was determined by hand harvesting two 5-m strips from two central rows of each plot. Sub-samples of seed cotton were ginned on a 20 saw Centennial Gin and lint samples were sent to the Fiber & Biopolymer Research Institute (Lubbock, Texas, USA) for fiber quality analysis using the High Volume Instrument (HVI) system. The following fiber properties were measured: micronaire (MIC), fiber length (mm), uniformity index (%), fiber strength (g tex^-1^), elongation, and color parameters, including reflectance (Rd), which indicates fiber brightness, and yellowness (+b).

### 2.4. Data analysis

All statistical analyses were conducted considering a split plot design. Data were analyzed using Minitab Statistical Software 22 (Minitab, LLC, State College, PA, USA).

For parameters measured repeatedly over time—namely LWP, RWC, F_v_/F_m_, and fluorescence-based RLC parameters—linear mixed-effects models were applied, with variety, irrigation, and time of day included as fixed effects and day included as a random effect. LAI was analyzed using the same modelling framework, excluding time of day, as measurements were performed once per day. For leaf gas exchange parameters and difference between canopy and air temperature (ΔT), analyses were performed separately for each sampling day, as measurement times differed among days and were not consistent across the experimental period. For δ^13^C, total N and C/N ratio, analyses were also performed separately for each sampling day to detect differences among genotypes.

Significance of fixed effects was assessed using ANOVA, and post hoc comparisons were performed using Tukey’s test at p = 0.05. Prior to analysis, data were tested for normality using the Ryan–Joiner test and for homogeneity of variances using Levene’s test (α = 0.05). Pearson correlation coefficients were calculated to assess relationships among selected variables.

## 3. Results

### 3.1. Soil water content

The temporal trend of soil water content (SWC) for each combination of variety (NG 4190 and ST 4990) and irrigation level (FI and DI) at six soil depths (0–100 cm) is shown in Figure 2. SWC decreased with increasing soil depth throughout the season, with values ranging from 0.161 to 0.302 m^3^ m^-3^ along the profile.

**Figure 2.**
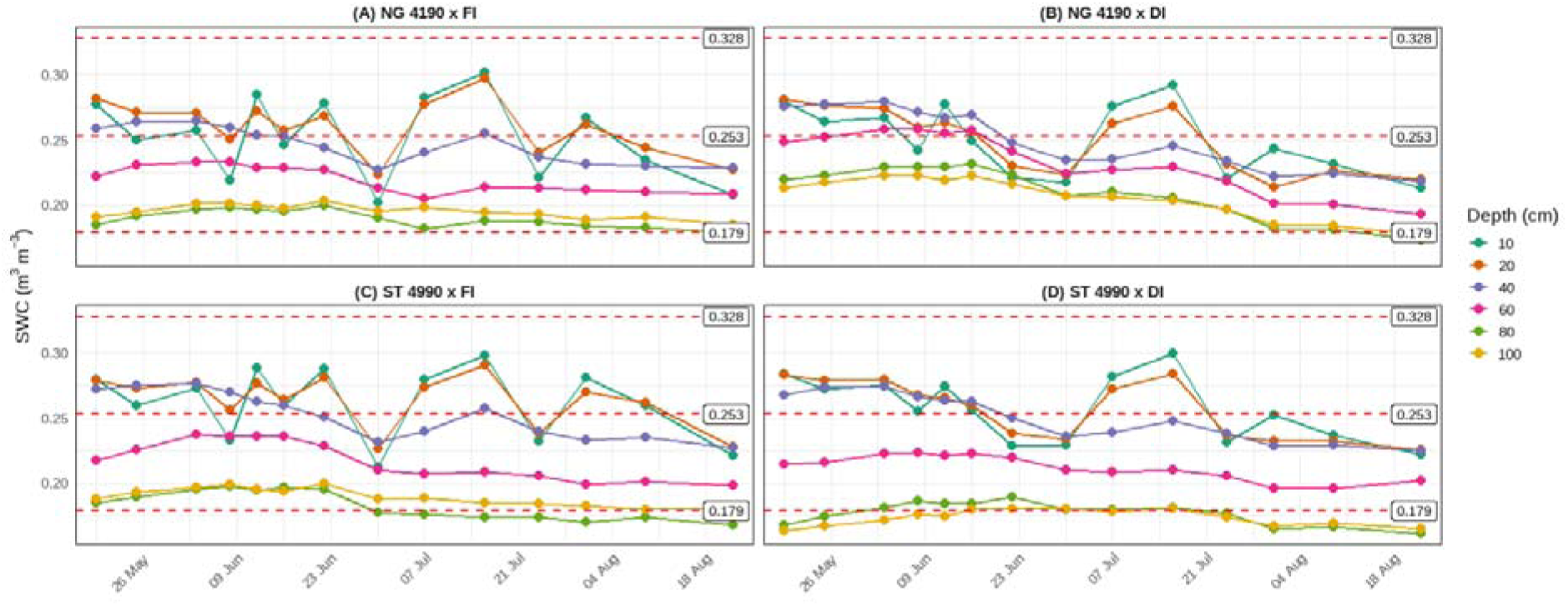
Soil water content over time for each combination of variety (NG 4190 and ST 4990) and irrigation regime (FI and DI) at six soil depths (0–100 cm). SWC = Soil water content; FI = full irrigation; DI = deficit irrigation. θ_fc_: 0.328 (m^3^ m^-3^); θ_wp_: 0.179 (m^3^ m^-3^); θ_p_: 0.253 (m^3^ m^-3^).

In the upper soil layers (0–20 cm), seasonal mean SWC values were approximately 0.26 m^3^ m^-3^, and remained above the threshold soil water content (0.253 m^3^ m^-3^) for most of the season. At 40 cm depth, SWC values exceeded this threshold only until late June (∼23 June), after which lower values were recorded. Lower SWC values were consistently observed at 60–100 cm compared to the upper layers (Figure 2A-D).

Differences between irrigation treatments were more evident in the deeper soil layers of ST 4990, where the SWC values were lower under DI (Figure 2D). In contrast, in NG 4190, SWC under deficit irrigation was slightly higher than under FI at 80–100 cm until mid-July, with mean differences of 0.02–0.03 m^3^ m^-3^ (Figure 2A-B). These differences diminished later in the season. Conversely, in the upper 20 cm, deficit irrigation generally resulted in lower SWC compared to full irrigation in both varieties during the late-season period.

### 3.2. Plant water status

Analysis of variance (ANOVA; Table 2) revealed a significant three-way interaction among variety, irrigation regime, and time of day on leaf water potential (LWP). In contrast, relative water content (RWC) was significantly affected by a two-way interaction.

**Table 2.**
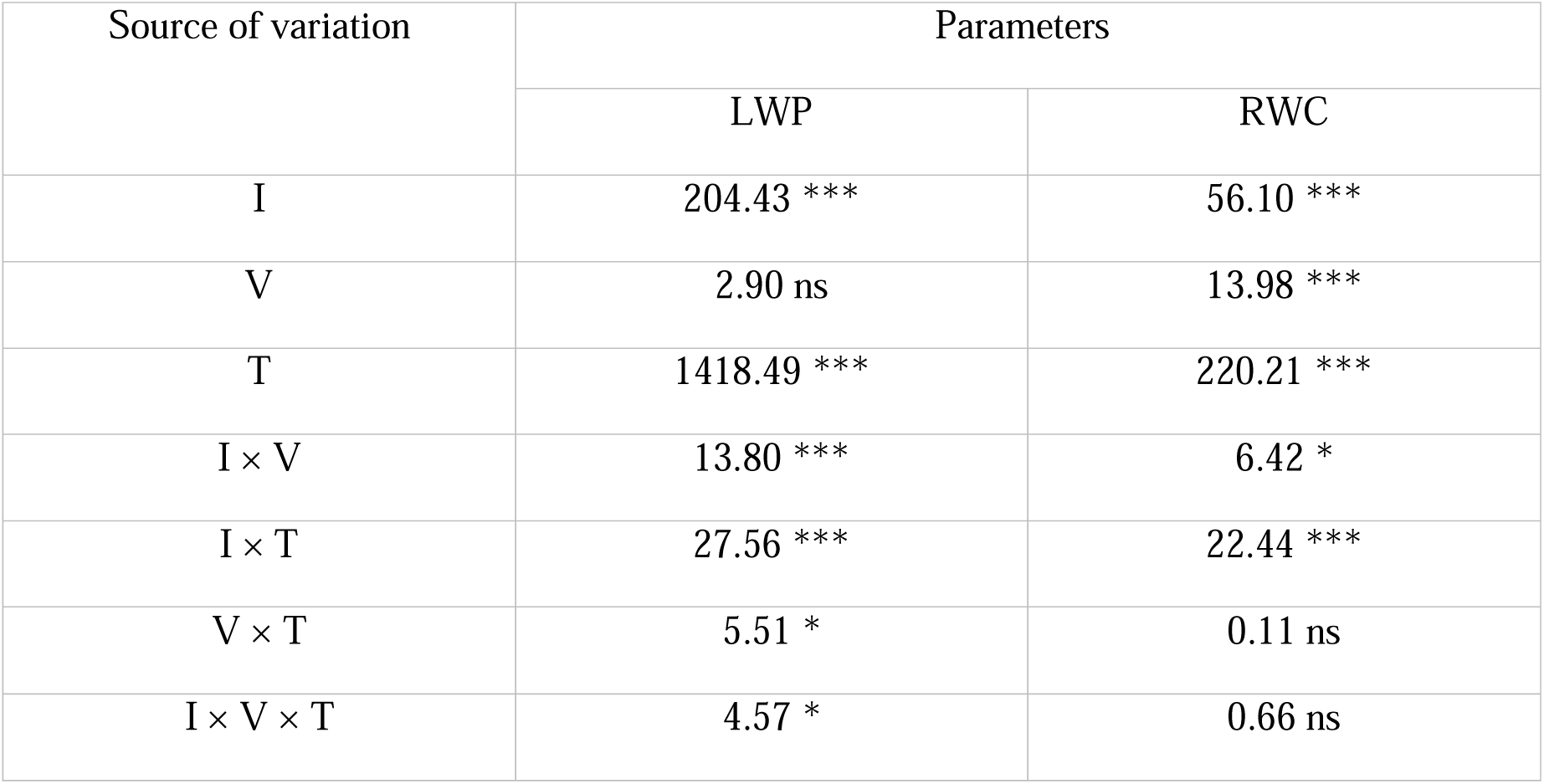
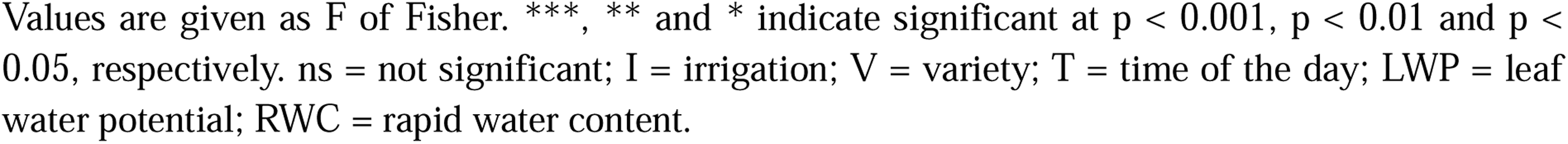
Analysis of variance (ANOVA) of main effects and interactions on LWP, RWC.

| Source of variation | Parameters |  |
| --- | --- | --- |
|  | LWP | RWC |
| I | 204.43 *** | 56.10 *** |
| V | 2.90 ns | 13.98 *** |
| T | 1418.49 *** | 220.21 *** |
| I × V | 13.80 *** | 6.42 * |
| I × T | 27.56 *** | 22.44 *** |
| V × T | 5.51 * | 0.11 ns |
| I × V × T | 4.57 * | 0.66 ns |

Values are given as F of Fisher. \*\*\*, \*\* and \* indicate significant at $p < 0.001$ , $p < 0.01$ and $p < 0.05$ , respectively. ns = not significant; I = irrigation; V = variety; T = time of the day; LWP = leaf water potential; RWC = rapid water content.

The variety × irrigation interaction at the two sampling times is shown in Fig. 3. At predawn, LWP ranged from −0.5 to −0.9 MPa, whereas midday values were more negative, ranging from −1.5 to −2.4 MPa. In both cases, the deficit irrigation treatment (DI) resulted in significantly more negative LWP compared with well-watered conditions (FI). Varietal differences were observed only at midday under full irrigation, with ST 4990 exhibiting LWP values approximately 0.3 MPa less negative than NG 4190.

**Figure 3.**
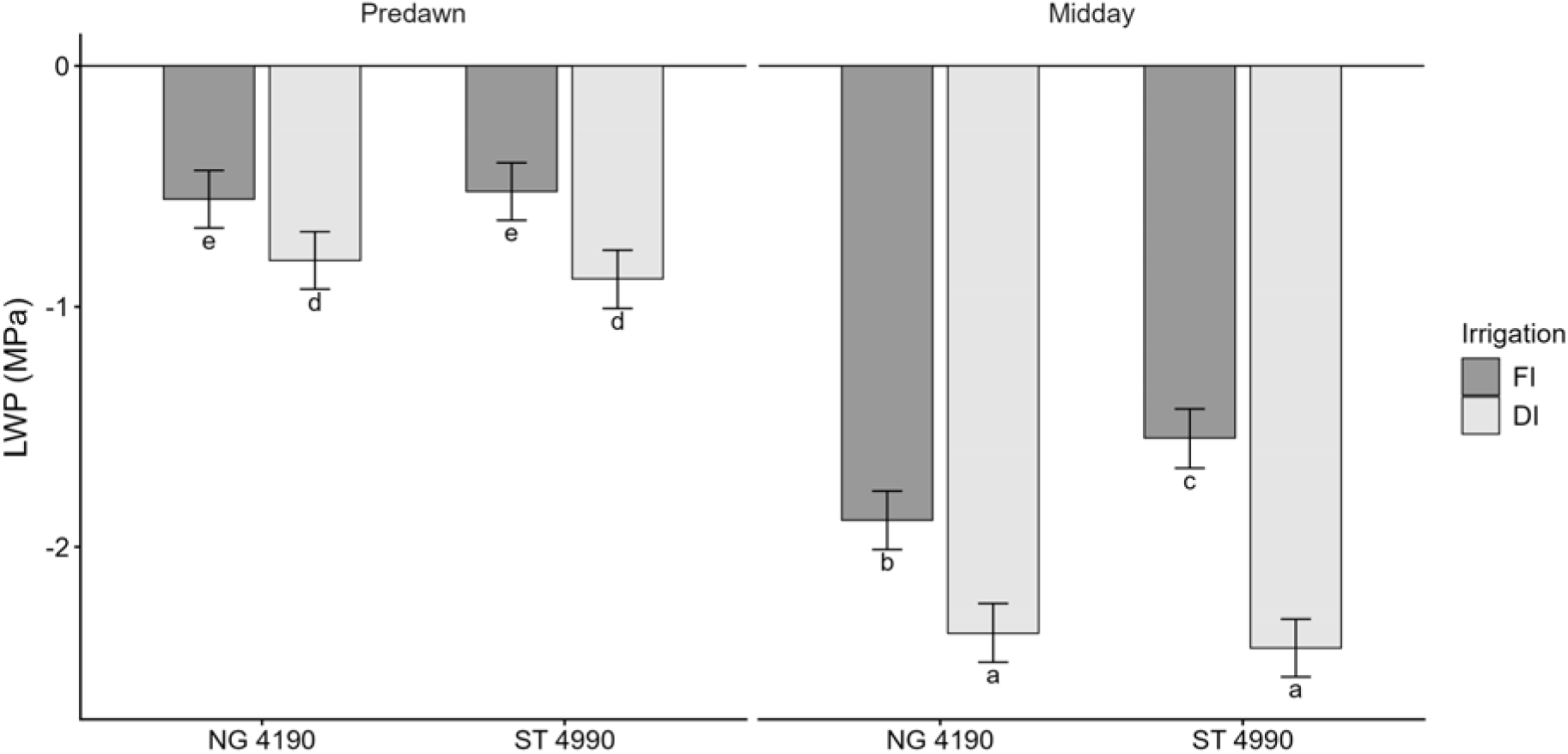
Effect of variety, irrigation regime, and time of the day (predawn; midday) on leaf water potential (LWP). Different letters indicate significant differences among means (p ≤ 0.05; Tukey’s test). Error bars represent the standard error (SE). Each treatment was replicated three times. Measurements were taken at predawn and midday across five days (D1–D5). Varieties were NG 4190 B3XF and ST 4990 B3XF; irrigation regimes were FI (full irrigation) and DI (deficit irrigation).

RWC ranged from 71.3% to 82.4% (Fig. 4), with the highest values recorded at predawn. ANOVA revealed that differences between treatments were significant at midday, where water-stressed plants exhibited the lowest RWC (Fig. 4A). Furthermore, under deficit irrigation (DI), ST 4990 showed a significantly lower RWC compared to NG 4190, which did not differ between the two irrigation treatments (Fig. 4B).

**Figure 4.**
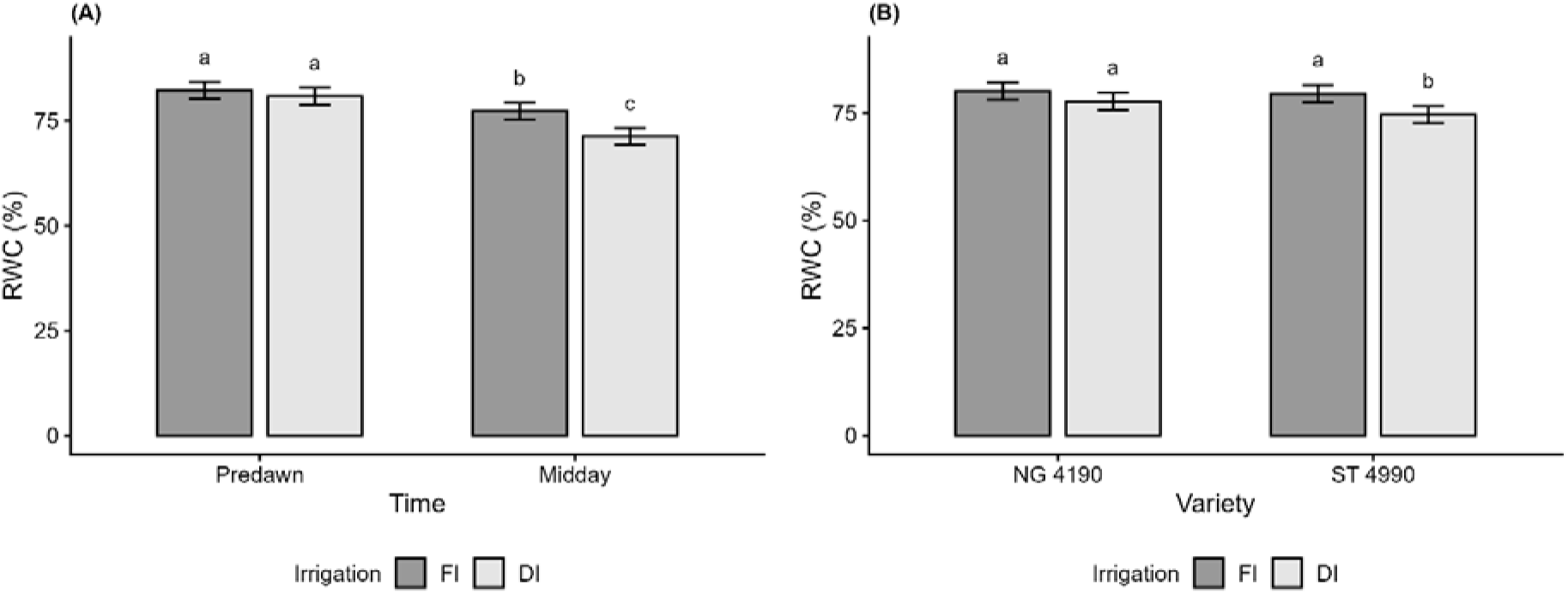
Effect of variety, irrigation regime, and time of the day (predawn; midday) on leaf water potential on relative water content (RWC). (A): irrigation × time; (B): variety × irrigation. Different letters indicate significant differences among means (p ≤ 0.05; Tukey’s test). Error bars represent the standard error (SE). Each treatment was replicated three times. Measurements were taken at predawn and midday across five days (D1–D5). Varieties were NG 4190 B3XF and ST 4990 B3XF; irrigation regimes were FI (full irrigation) and DI (deficit irrigation).

Regarding leaf area index (LAI), the mixed model analysis revealed that irrigation regime was the only significant factor affecting this trait, while variety and the variety × irrigation interaction were not significant (Table 3). Across the experimental season, deficit irrigation reduced LAI by approximately 30% compared with full irrigation (1.70 vs. 2.36, respectively; data not shown).

**Table 3.**
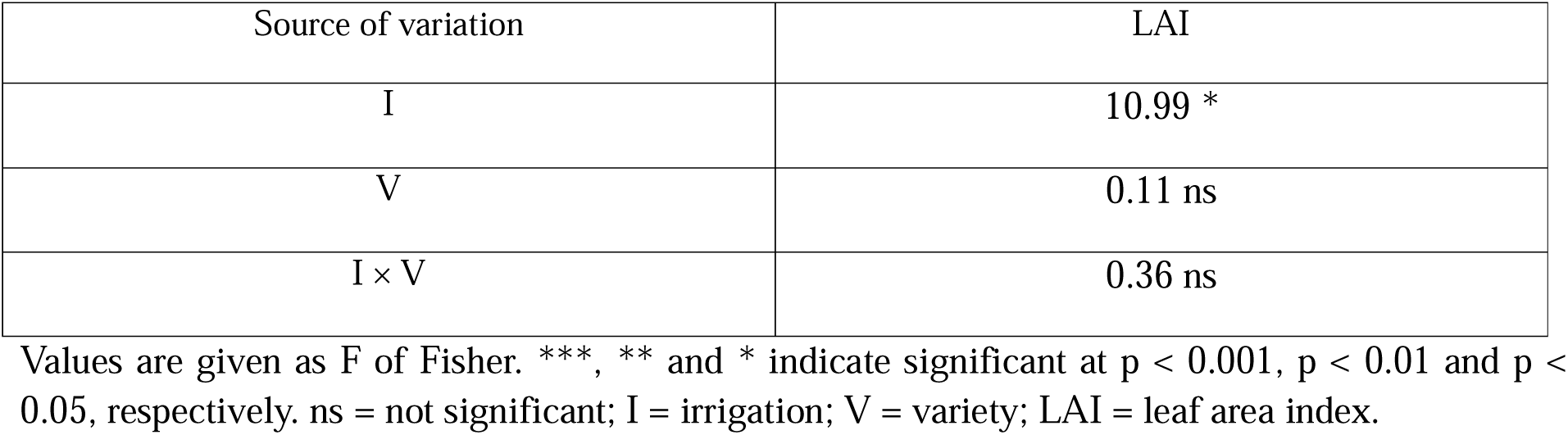
Analysis of variance (ANOVA) of main effects and interactions on LAI.

| Source of variation | LAI |
| --- | --- |
| I | 10.99 * |
| V | 0.11 ns |
| I × V | 0.36 ns |

Values are given as F of Fisher. \*\*\*, \*\* and \* indicate significant at $p < 0.001$ , $p < 0.01$ and $p < 0.05$ , respectively. ns = not significant; I = irrigation; V = variety; LAI = leaf area index.

### 3.3. Physiological responses

Analysis of gas exchange responses in the two cotton varieties under different irrigation regimes revealed differentiated responses among the measured parameters, both under steady-state CO□ and PPFD conditions and during measurements conducted under varying CO□ and PPFD (Tables 4 and 4).

**Table 4.** Analysis of variance (ANOVA) of main effects and interactions on instantaneous gas exchange parameters (A, g_s_, and E), and ΔT.

|  | Parameters | Source of variation |  |  |  |  |  |  |
| --- | --- | --- | --- | --- | --- | --- | --- | --- |
| | | I | V | T | $I \times V$ | $I \times T$ | $V \times T$ | $I \times V \times T$ |

|  |  |  |  |  |  |  |  | T |
| --- | --- | --- | --- | --- | --- | --- | --- | --- |
| D1 | A | 21.98 ns | 0.06 ns | 163.41<br>*** | 0.02 ns | 47.32 * | 27.67<br>ns | 0.15 ns |
|  | g<sub>s</sub> | 20.16 ns | 0.03 ns | 270.52<br>*** | 0.26 ns | 130.02<br>*** | 44.13<br>* | 0.25 ns |
|  | E | 35.84 ns | 0.02 ns | 264.22<br>*** | 0.04 ns | 37.05 * | 25.14<br>ns | 0.09 ns |
|  | ΔT | 0.54 ns | 0.46 ns | 242.28<br>*** | 64.75<br>* | 116.88<br>*** | 0.03 ns | 0.83 ns |
| D2 | A | 594.90<br>*** | 0.41 ns | 479.60<br>*** | 10.41<br>ns | 151.20<br>*** | 21.08<br>ns | 0.66 ns |
|  | g<sub>s</sub> | 136.98<br>*** | 0.01 ns | 341.46<br>*** | 16.87<br>ns | 267.01<br>*** | 51.44<br>** | 31.61 * |
|  | E | 580.19<br>*** | 0.18 ns | 330.76<br>*** | 13.35<br>ns | 283.30<br>*** | 33.85<br>* | 0.98 ns |
|  | ΔT | 189.76<br>*** | 0.01 ns | 110.40<br>*** | 15.06<br>ns | 901.81<br>*** | 0.42 ns | 27.19<br>ns |
| D3 | A | 0.79 ns | 0.00 ns | 487.36<br>*** | 0.09 ns | 40.78 * | 0.40 ns | 0.96 ns |
|  | g<sub>s</sub> | 19.43 ns | 10.24 ns | 828.71<br>*** | 0.03 ns | 71.37 ** | 0.08 ns | 0.22 ns |
|  | E | 0.42 ns | 0.73 ns | 362.99<br>*** | 0.12 ns | 53.05 ** | 0.36 ns | 0.23 ns |
|  | ΔT | 0.11 ns | 0.58 ns | 21.48 ns | 0.27 ns | 0.17 ns | 16.98<br>ns | 21.32<br>ns |
| D4 | A | 548.99<br>*** | 0.00 ns | 153.85<br>*** | 0.00 ns | 639.64<br>*** | 0.51 ns | 23.47<br>ns |
|  | g<sub>s</sub> | 887.43<br>*** | 132.04<br>*** | 754.88<br>*** | 17.90<br>ns | 318.01<br>*** | 11.34<br>ns | 0.26 ns |
|  | E | 936.70<br>*** | 58.69 * | 178.85<br>*** | 0.11 ns | 805.96<br>*** | 0.03 ns | 0.07 ns |
|  | ΔT | 136.60<br>*** | 13.18 ns | 313.69<br>*** | 0.75 ns | 268.39<br>*** | 0.37 ns | 0.61 ns |
| D5 | A | 0.24 ns | 33.39 ns | 58.46 ** | 10.92<br>ns | 190.80<br>*** | 19.89<br>ns | 26.98<br>ns |
| | $g_s$ | 0.22 ns | 47.43 ns | 89.34<br>*** | 10.29<br>ns | 105.35<br>*** | 26.63<br>ns | 17.66<br>ns |
|  | E | 0.05 ns | 67.50 * | 504.13<br>*** | 0.68 ns | 198.05<br>*** | 0.77 ns | 23.02<br>ns |
| | $\Delta T$ | 225.12<br>*** | 0.23 ns | 129.78<br>*** | 0.35 ns | 660.33<br>*** | 0.38 ns | 16.78<br>ns |

Values are given as F of Fisher. \*\*\*, \*\* and \* indicate significant at $p < 0.001$ , $p < 0.01$ and $p < 0.05$ , respectively. ns = not significant; I = irrigation; V = variety; T = time of the day; D = day (D1 = 9 June, D2 = 25 June, D3 = 10 July, D4 = 1 August, D5 = 28 August); A = net photosynthesis rate; $g_s$ = stomatal conductance; E = transpiration rate; $\Delta T$ = difference between canopy and air temperature.

Instantaneous gas exchange parameters, including net photosynthesis (A), stomatal conductance (g_s_), and transpiration (E), showed statistically significant I × T interactions, with occasional significant effects for V × T. No significant effect of the I × V interaction was observed. Although the main focus of this study was to use physiological indices to select cotton varieties with higher drought tolerance, the significant interaction I × T is discussed here, as the analysis of diurnal physiological patterns provides additional insight into plant responses to the two irrigation regimes. Rather than presenting a separate figure specifically for the I × T interaction, Figure 5 reports the diurnal trends across the measurement periods for each variety × irrigation combination over the five measurement days (D1–D5). This representation was considered more informative and facilitates interpretation, while still allowing clear visualization of the irrigation × time-of-day interaction.

**Figure 5.**
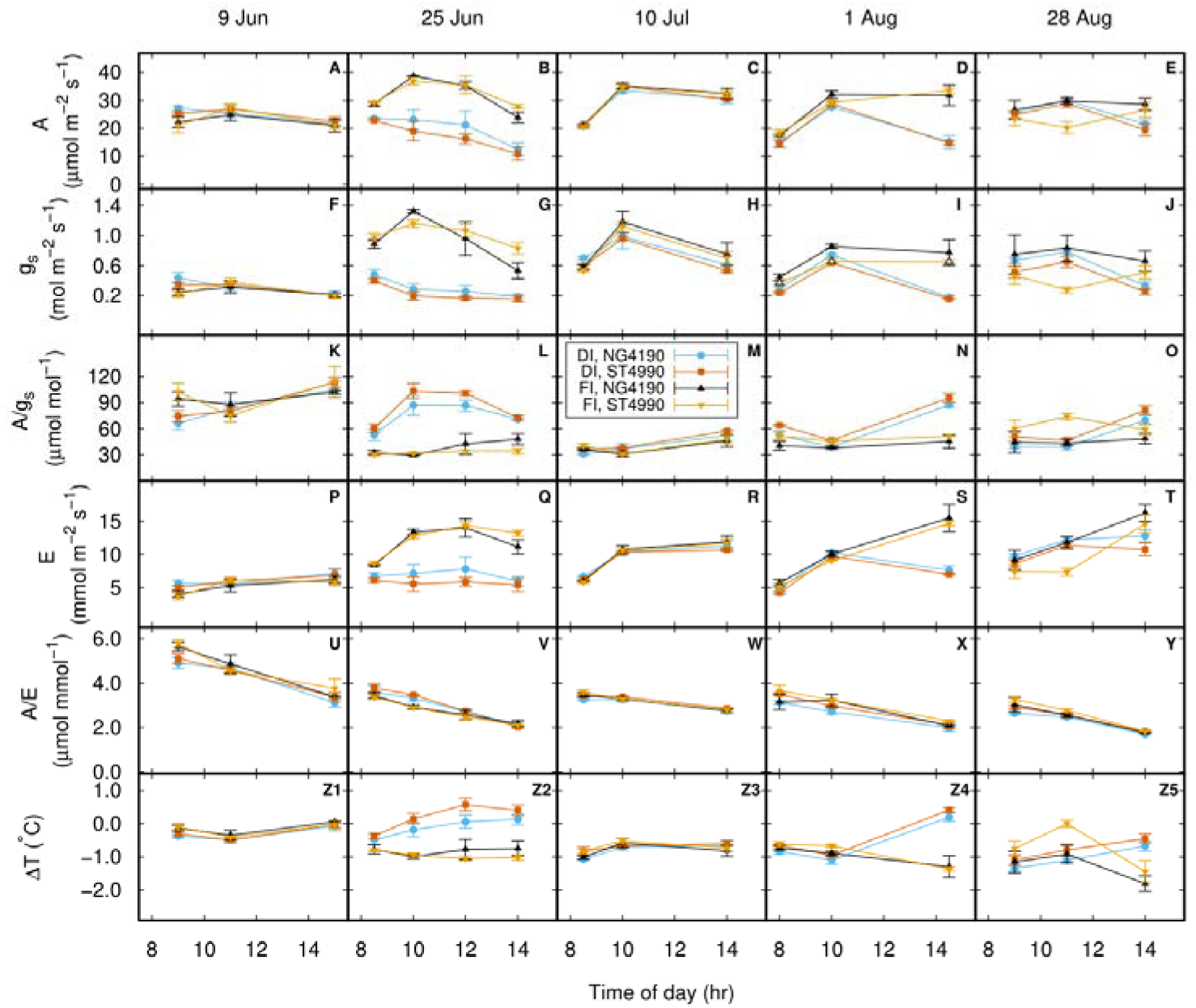
Diurnal courses of net photosynthetic rate (A shown in panels A through E), stomatal conductance (g_s_, panels F through J), intrinsic water use efficiency (A/g_s_, panels K through O), transpiration rate (E, panels P through T), instantaneous water use efficiency (A/E, panels U through Y), and leaf temperature minus air temperature (ΔT, panels Z1 through Z5) of two cotton varieties, i.e., NG 4190 and ST 4990, measured over five clear days in summer 2025 at Uvalde under full irrigation (FI) and deficit irrigation (DI) conditions. On each day, the measurement was made 3-4 times from 8:00 am to 15:00 pm local time, on three randomly chosen sunlit mature leaves from each of the treatments (n = 3, error bars indicate ± one standard errors of the means). The gas exchange measurement was made under field conditions with LI-6400XT Portable Photosynthesis System, and the canopy temperature was measured using an infrared radiometer (Model MI-210). The five days chosen included 9 June, 25 June, 10 July, 1 August, and 28 August, 2025.

The only instance in which a significant I × V × T interaction was observed occurred on the second measurement date (D2 = 25 June) for stomatal conductance (g_s_), coinciding with a more pronounced water stress in plants subjected to deficit irrigation (DI), as also indicated by the increase in ΔT (see below). On this date, A under DI showed an average reduction of approximately 40% compared with FI, while transpiration was significantly higher under well-watered conditions. Under deficit irrigation, ST 4990 consistently exhibited lower g_s_ values than NG 4190 across all measurement times, although differences were not always statistically significant (data not shown). Stomatal conductance (0.17-1.15 mol m^-2^ s^-1^) generally reached higher values at the second measurement time (∼11:00), with differences between FI and DI not always statistically significant across all measurement times (Fig. 5F-J). A ranged from 17.9 to 37.9 µmol m^-2^ s^-1^ under FI and from 11.8 to 34.3 µmol m^-2^ s^-1^ under DI, with statistically significant differences between the two irrigation regimes observed mainly in the early afternoon (∼14:00) (Fig. 5A-E). Similarly, for E, the most pronounced differences between irrigation regimes were observed at the last measurement time, when well-watered plants maintained higher transpiration rates compared with those under deficit irrigation (Fig. 5P-T)).

Intrinsic water use efficiency (A/g_s_) tended to be higher under FI compared to DI during the more stressful days (D2, D4, D5), with peak values observed during the hottest hours of the day (Fig. 5K-O). Instantaneous water use efficiency (A/E) generally declined throughout the day in all treatments, reflecting the diurnal pattern of transpiration and photosynthesis (Fig. 5U-Y).

The canopy-to-air temperature difference (ΔT) was found to be highly significant only for the I × T interaction, except for the third measurement (D3 = 10 July), which did not show statistically significant differences at any level (Table 4). Similarly, the daily trends of ΔT for each variety × irrigation combination are reported in Figure 5Z1-Z5 to facilitate interpretation of the I × T interaction. In plants under FI, ΔT ranged between −1.4 and 1.9 °C throughout the measurement period, with a single peak of 5.4 °C recorded at 14:30 on the first day (D1 = 9 June). Under DI, values ranged from 0.7 to 7.1 °C, with the most pronounced differences observed during third measurement time (∼14:00), while the second measurement date reflected the most evident stress. Throughout the day, under water deficit, ΔT increased in parallel with rising temperatures, whereas under full irrigation, canopy temperature remained moderated even during the hottest hours.

Photosynthetic parameters estimated from A/C_i_ curves, including the maximum Rubisco carboxylation rate (V_cmax_), the maximum electron transport rate (J_max_), and dark respiration (R_d_), did not show statistically significant differences among treatments. Instead, triose phosphate utilization (TPU) was significantly affected, with the lowest values observed in NG 4190 under deficit irrigation (Table 5).

**Table 5.** Analysis of variance (ANOVA) of interactions on biochemical, photochemical, and stomatal parameters derived from A/C□ curve fitting, light response curve analysis, and the Medlyn stomatal conductance model.

| Parameters | NG 4190 ×<br>FI | NG 4190 ×<br>DI | ST 4990 ×<br>FI | ST 4990 ×<br>DI | Significance |
| --- | --- | --- | --- | --- | --- |
| $V_{\text{cmax}}$ ($\mu\text{mol CO}_2 \text{ m}^{-2} \text{ s}^{-1}$) | 108.9±11.2 | 88.1±11.9 | 105.4±11.0 | 78.9±14.6 | ns |
| $J_{\max}$ ($\mu\text{mol CO}_2 \text{ m}^{-2} \text{ s}^{-1}$) | 212.5±19.8 | 129.8±17.3 | 204.9±12.1 | 99.2±25.6 | ns |
| $R_d$ ($\mu\text{mol CO}_2 \text{ m}^{-2} \text{ s}^{-1}$) | 2.038±0.304 | 2.068±0.128 | 2.143±0.186 | 1.481±0.143 | ns |
| TPU ($\mu\text{mol CO}_2 \text{ m}^{-2} \text{ s}^{-1}$) | 14.56±1.23 a | 8.69±0.52 b | 14.27±1.03 a | 10.64±2.42 ab | * |
| $P_m$ ($\mu\text{mol CO}_2 \text{ m}^{-2} \text{ s}^{-1}$) | 54.68±4.43 a | 38.15±3.93 ab | 47.59±2.91 a | 27.69±5.12 b | * |
| $\alpha$ | 0.0813±0.0014 ab | 0.0790±0.0032 ab | 0.0860±0.0009 a | 0.0603±0.0104 b | * |
| $\theta$ | 0.2575±0.0932 | 0.2197±0.1848 | 0.4138±0.0705 | 0.6783±0.0996 | ns |
| $g_1$ | 7.259±0.169 a | 5.99± b | 6.702±0.162 a | 5.302±0.142 c | * |

Values represent means ± standard error (SE). Different letters indicate significant differences among means ( $p \leq 0.05$ ; Tukey's test). \*\*\*, \*\* and \* indicate significance at $p < 0.001$ , $p < 0.01$ and $p < 0.05$ , respectively; ns = not significant. Each treatment was replicated at least three times. The values of the parameter $g_1$ were compared via ANOVA in GraphPad Prism according to the nonlinear regression results (mean, SE and $n=144$ ; see Supplementary material). Varieties were NG 4190 B3XF and ST 4990 B3XF; irrigation regimes were FI (full irrigation) and DI (deficit irrigation). $V_{\max}$ = maximum carboxylation rate of rubisco; $J_{\max}$ = maximum electron transport rate; $R_d$ = dark respiration rate; TPU = triose phosphate utilization; $P_m$ = maximum photosynthesis; $\alpha$ = photosynthesis efficiency; $\theta$ = photosynthesis sharpness parameter; $g_1$ = the Medlyn stomatal optimization parameter.

In contrast, parameters derived from light-response curves revealed a significant effect of the I × V interaction (Table 5). Specifically, the maximum photosynthesis under high irradiance (P_m_) and the photosynthetic efficiency at low light intensities (α) showed statistically significant differences between treatments. P_m_ ranged from 27.69 to 54.68 μmol CO□ m^-2^ s^-1^, recorded in ST 4990 × DI and NG 4190 × FI, respectively. While no significant differences were observed between the two cotton varieties under full irrigation, under deficit irrigation, ST 4990 showed a ∼27% reduction in P_m_ compared with NG 4190. A similar trend was observed for photosynthetic efficiency (α), with the lowest value recorded in ST 4990 under deficit irrigation (0.06). By contrast, the curvature parameter of the light-response curve (θ) did not differ significantly among treatments.

Figure 6 shows that the light-response curves were largely overlapping between the two varieties under full irrigation (FI), whereas under deficit irrigation (DI) they exhibited a clear separation across the entire PPFD gradient, with more pronounced differences at high light intensities. Additionally, photosynthesis tended to reach marginal increases at PPFD values above ∼1000–1200 *µ*mol m^-2^ s^-1^ under FI, while under DI, the light-response was already limited at PPFD values around 500–700 *µ*mol m^-2^ s^-1^.

**Figure 6.**
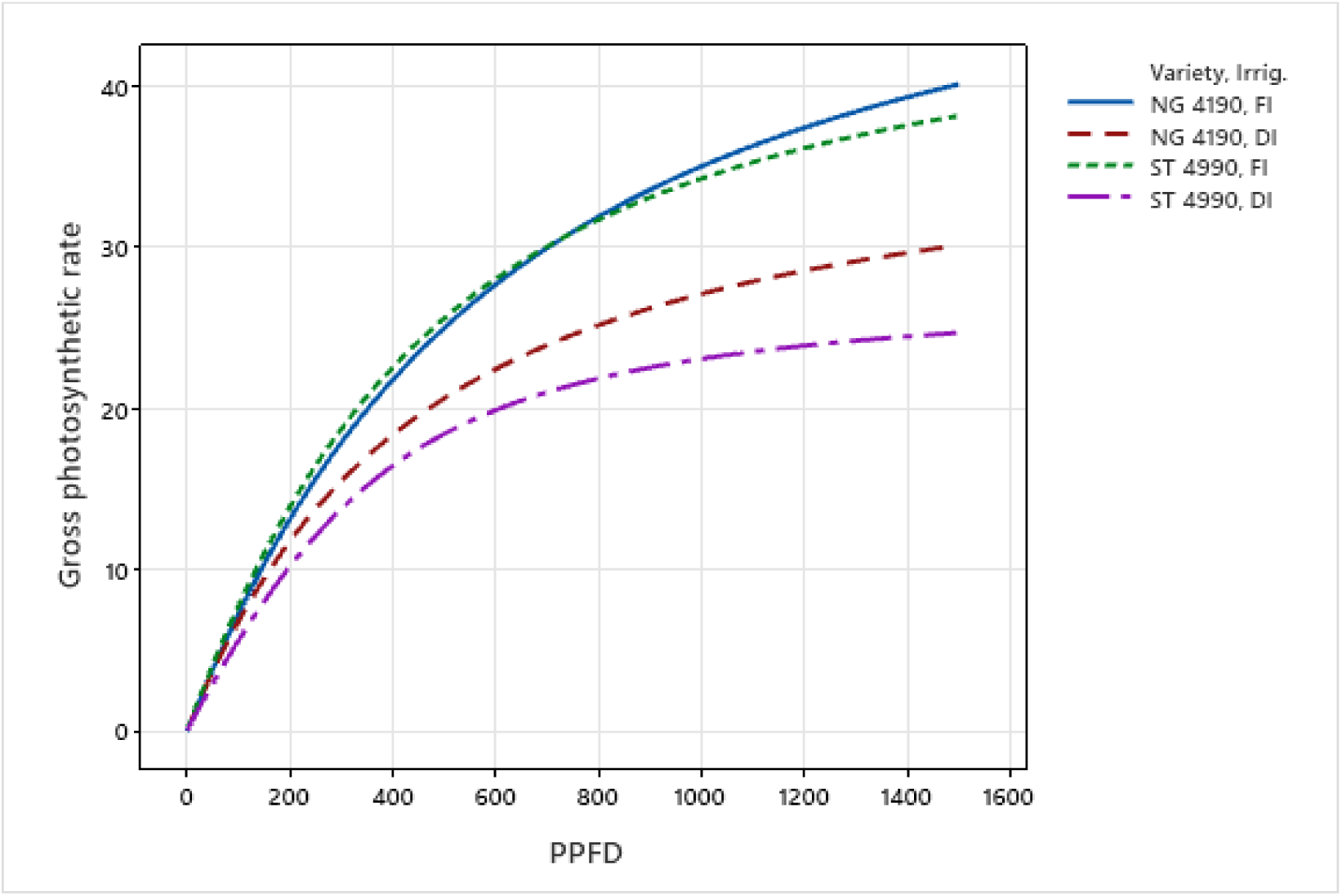
Photosynthesis–light response curves of two cotton varieties under contrasting irrigation regimes, fitted with a non-rectangular hyperbola model using mean parameter estimates (α, P_m_, θ) from 3–4 replicates per treatment. Values of the fitted parameters are shown in Table 5.

Finally, the stomatal optimization parameter (g□, Table 5) ranged from 5.3 to 7.3. The highest values were observed under full irrigation, where no statistically significant differences were found between the two varieties. In contrast, under deficit irrigation, ST 4990 exhibited a significantly lower g□ value compared with NG 4190 (5.30 vs. 5.99). Consistently, regression analysis (Fig. 7) between g_s_ and the photosynthetic demand normalized by atmospheric CO□ concentration and vapor pressure deficit (A/(Ca√D)) showed that, under deficit irrigation, NG 4190 exhibited a significantly steeper slope than ST 4990, indicating greater stomatal conductance per unit of photosynthetic demand.

**Figure 7.**
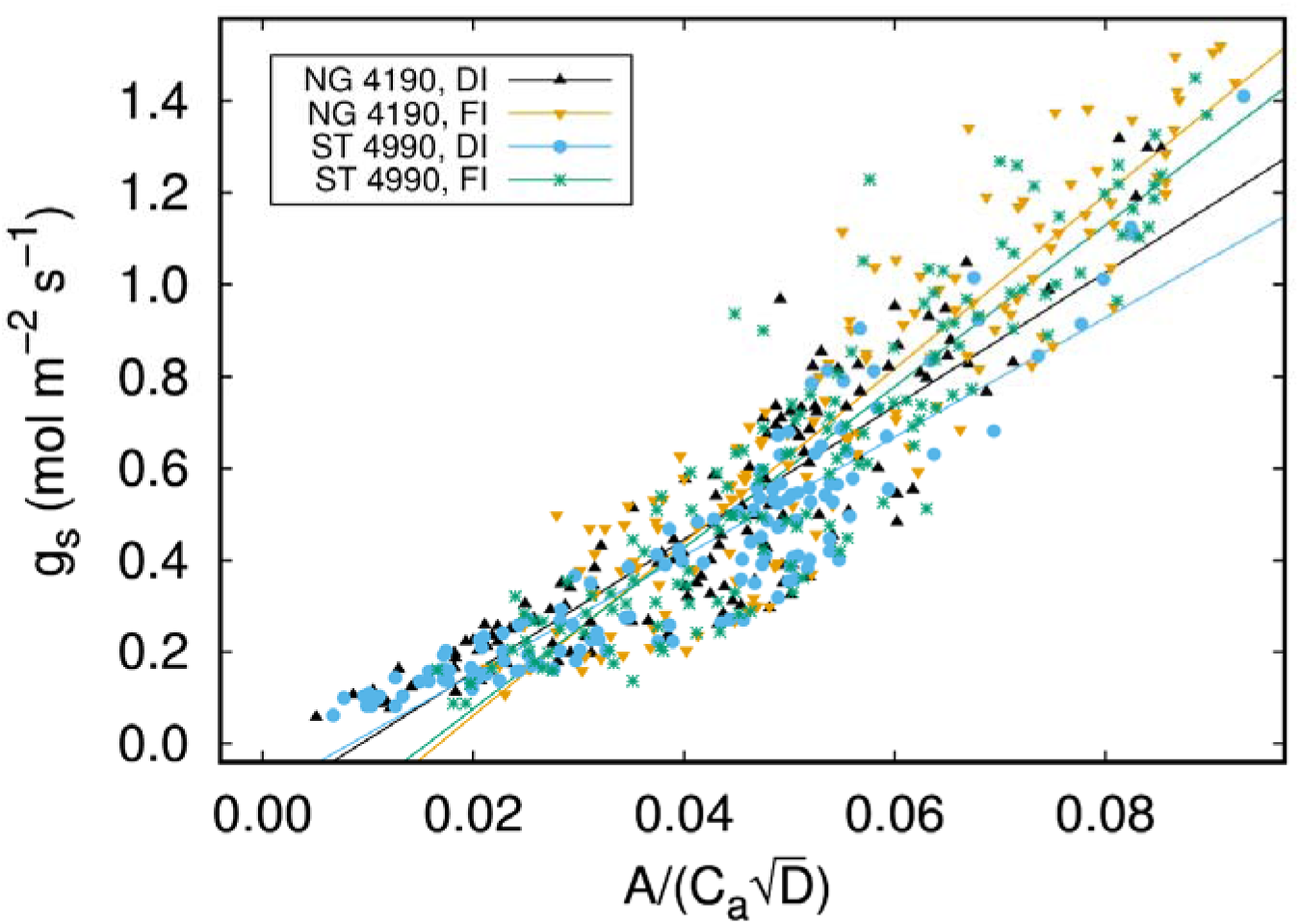
Visualization of stomatal behavior for two cotton varieties by regressing leaf stomatal conductance (g_s_, mol m^-2^ s^-1^) against 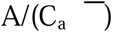 measured under full irrigation (FI) and deficit irrigation (DI) conditions. A is net photosynthetic rate (μmol m^-2^ s^-2^), C_a_ is the atmospheric CO_2_ concentration at the leaf surface (μmol mol^-1^), and D is leaf-to-air vapor pressure deficit (kPa). Each data point represents one set of values of 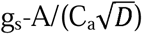 of a leaf of one cotton variety measured at a specific time on 9 June, 25 June, 10 July, 1 August, or 28 August, 2025 (all clear days) under field conditions. The slope of the regression line is inversely proportional to leaf water use efficiency.

Results of chlorophyll fluorescence parameters are summarized in Table 6. The photochemical efficiency of PSII (F_v_/F_m_) did not show a significant V × I interaction, but a statistically significant effect of variety was observed, with ST 4990 haivng a higher value than NG 4190 (0.839 vs. 0.823, Fig. 8A).

**Figure 8.**
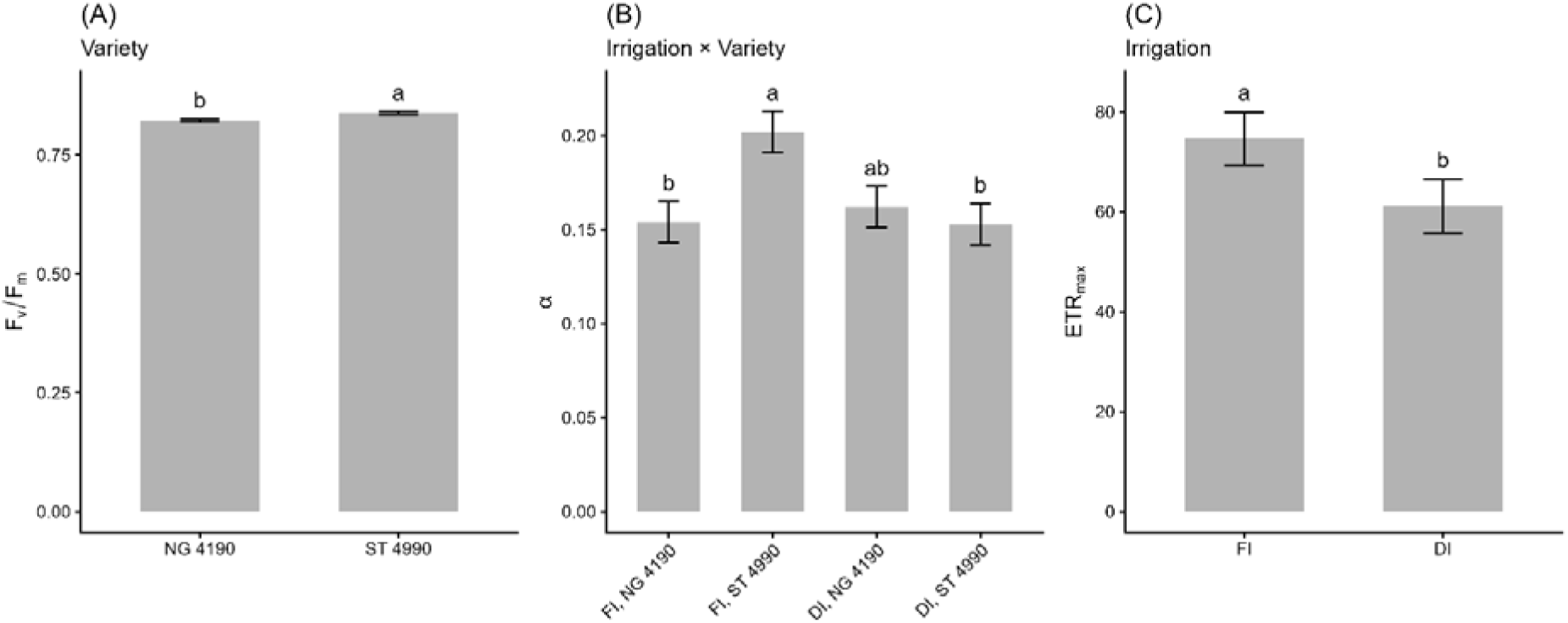
Chlorophyll fluorescence parameters in response to genotype, irrigation, and their interaction. (A) F_v_/F_m_, (B) α_PSII_, (C) ETR_max_. Different letters indicate significant differences among means (p ≤ 0.05; Tukey’s test). Error bars represent the standard error (SE). ***, ** and * indicate significance at p < 0.001, p < 0.01 and p < 0.05, respectively. Each treatment was replicated three times. Measurements of F_v_/F_m_ were taken on 1 and 28 August, and RLC on 9 and 28 August. Varieties were NG 4190 B3XF and ST 4990 B3XF; irrigation regimes were FI (full irrigation) and DI (deficit irrigation). F_v_/F_m_ = maximum quantum efficiency of PSII; α_PSII_ = initial slope of the rapid light curve (photosynthetic quantum efficiency); ETR_max_ = maximum electron transport rate.

**Table 6.**
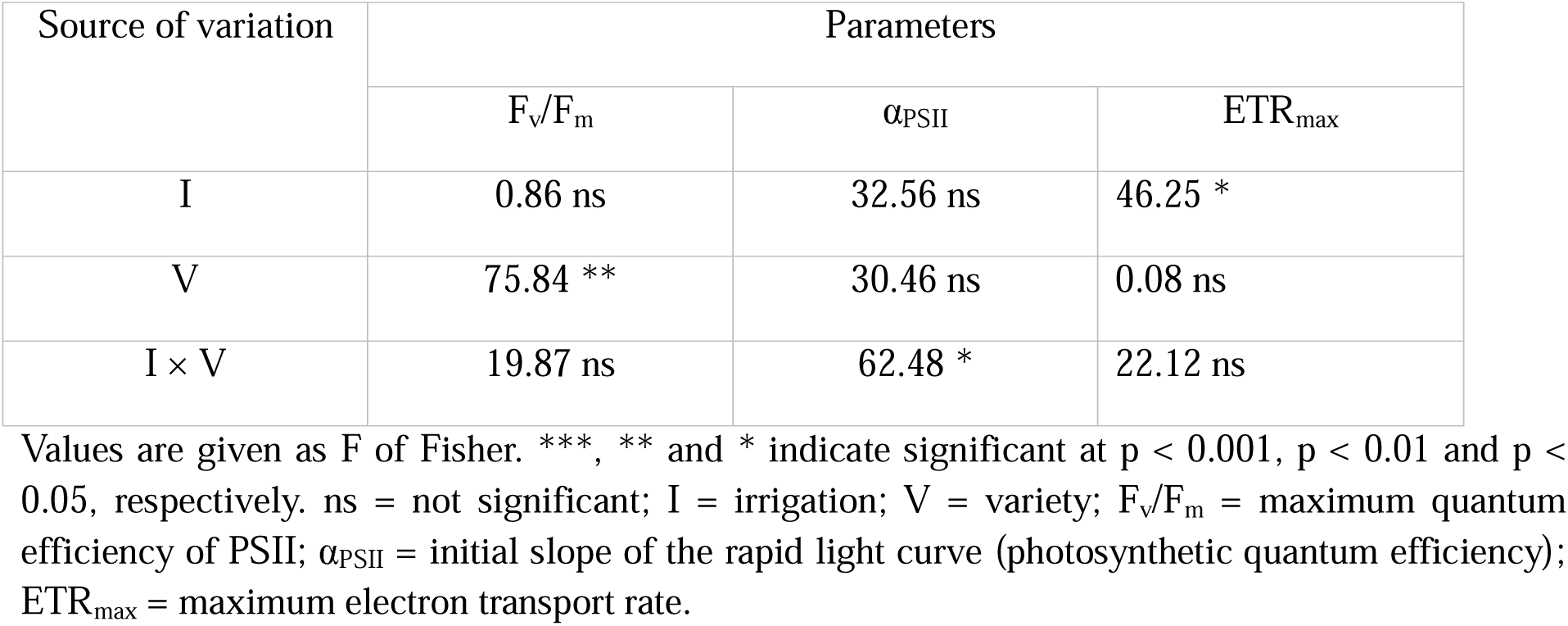
Analysis of variance (ANOVA) of main effects and interactions on chlorophyll fluorescence parameters.

| Source of variation | Parameters |  |  |
| --- | --- | --- | --- |
| | $F_v/F_m$ | $\alpha_{PSII}$ | $ETR_{max}$ |
| I | 0.86 ns | 32.56 ns | 46.25 * |
| V | 75.84 ** | 30.46 ns | 0.08 ns |
| $I \times V$ | 19.87 ns | 62.48 * | 22.12 ns |

Values are given as F of Fisher. \*\*\*, \*\* and \* indicate significant at $p < 0.001$ , $p < 0.01$ and $p < 0.05$ , respectively. ns = not significant; I = irrigation; V = variety; $F_v/F_m$ = maximum quantum efficiency of PSII; $\alpha_{PSII}$ = initial slope of the rapid light curve (photosynthetic quantum efficiency); $ETR_{max}$ = maximum electron transport rate.

For rapid light-response curve (RLC) parameters, the initial slope (α_PSII_), indicative of apparent quantum efficiency, was influenced by the V × I interaction, whereas the maximum electron transport rate (ETR_max_) was determined solely by the irrigation regime (Table 6). The highest α_PSII_ value was observed in ST 4990 under optimal irrigation, while under deficit irrigation, NG 4190 had a slightly higher α_PSII_ than ST 4990 (Fig. 8B). ETR_max_ was 18% higher in FI than in DI, highlighting the positive effect of full water availability on the maximum electron transport capacity of PSII (Fig. 8C).

### 3.4. Isotopic composition and leaf nitrogen status

The ANOVA performed on carbon isotope composition (δ^13^C), including day as a fixed factor, did not reveal significant effects, although the p-value for genotype was relatively low (p = 0.068; data not shown). However, when analyzed for each sampling day separately, a significant difference emerged only on the last sampling day (D5, August 28) for the variety factor (Table 7). Regarding TKN and C/N ratio, no significant effects were observed across the five sampling dates, except at the second sampling date (D2, June 25) (Table 7). Nevertheless, Tukey’s post hoc test for TKN at D2 did not identify significant pairwise differences among treatments.

**Table 7.**
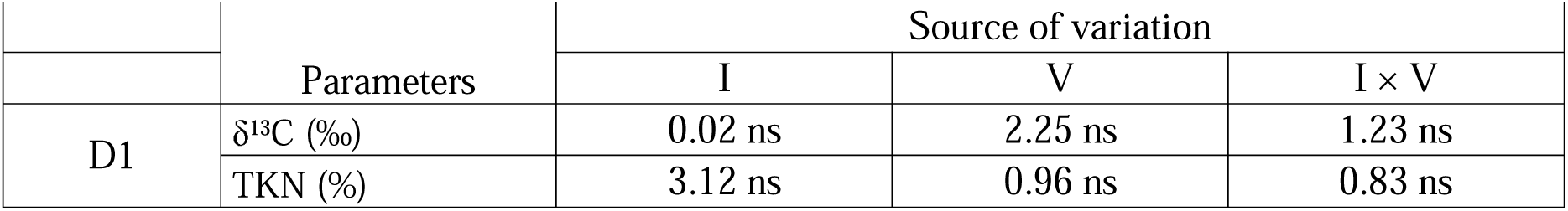
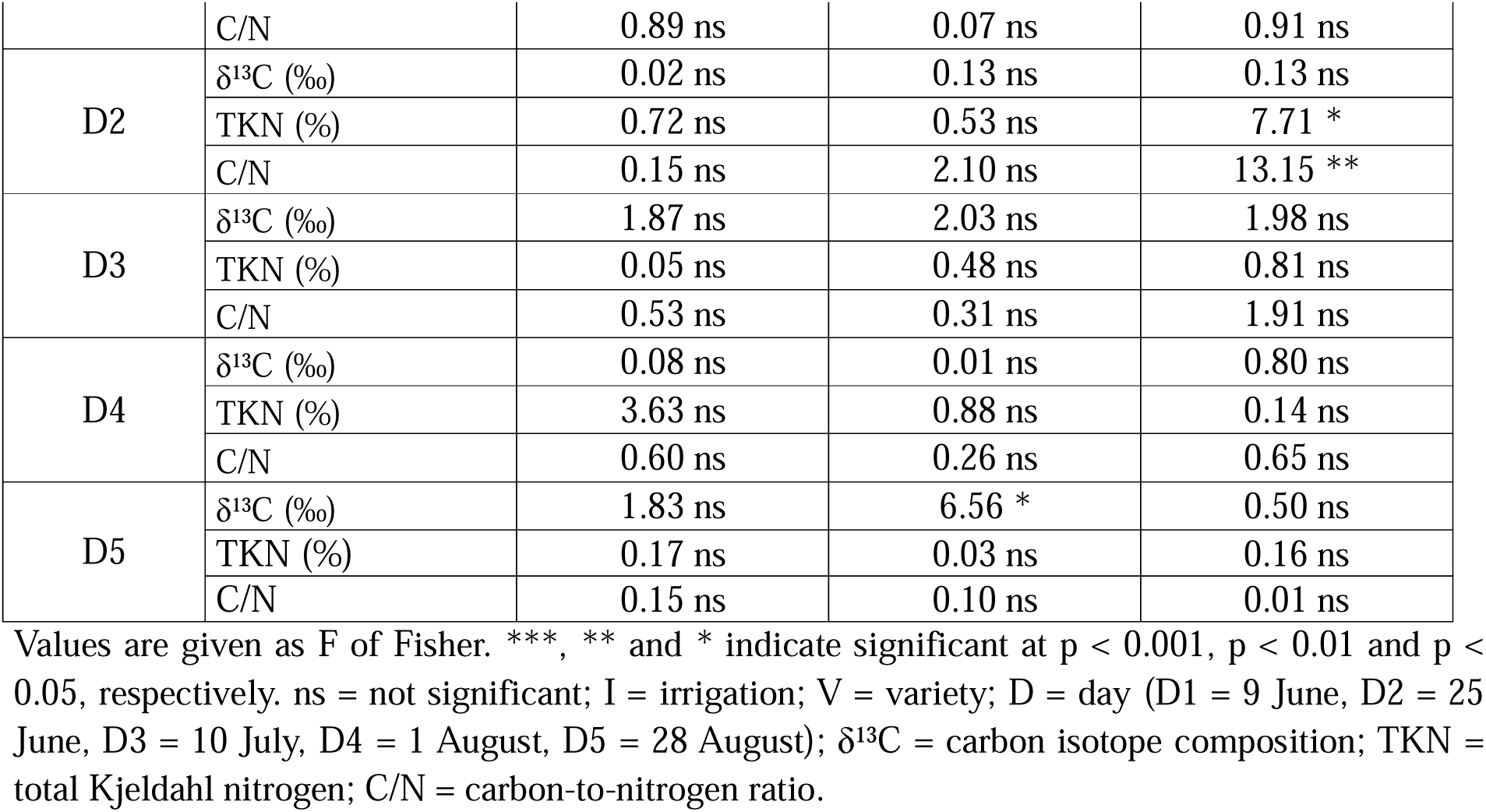
Analysis of variance (ANOVA) of main effects and interaction on carbon isotope composition, total nitrogen, and C/N ratio.

|  | Parameters | Source of variation |  |  |
| --- | --- | --- | --- | --- |
|  |  | I | V | I × V |
| D1 | $\delta^{13}C$ (‰) | 0.02 ns | 2.25 ns | 1.23 ns |
|  | TKN (%) | 3.12 ns | 0.96 ns | 0.83 ns |
|  | C/N | 0.89 ns | 0.07 ns | 0.91 ns |
| D2 | $\delta^{13}\text{C}$ (‰) | 0.02 ns | 0.13 ns | 0.13 ns |
|  | TKN (%) | 0.72 ns | 0.53 ns | 7.71 * |
|  | C/N | 0.15 ns | 2.10 ns | 13.15 ** |
| D3 | $\delta^{13}\text{C}$ (‰) | 1.87 ns | 2.03 ns | 1.98 ns |
|  | TKN (%) | 0.05 ns | 0.48 ns | 0.81 ns |
|  | C/N | 0.53 ns | 0.31 ns | 1.91 ns |
| D4 | $\delta^{13}\text{C}$ (‰) | 0.08 ns | 0.01 ns | 0.80 ns |
|  | TKN (%) | 3.63 ns | 0.88 ns | 0.14 ns |
|  | C/N | 0.60 ns | 0.26 ns | 0.65 ns |
| D5 | $\delta^{13}\text{C}$ (‰) | 1.83 ns | 6.56 * | 0.50 ns |
|  | TKN (%) | 0.17 ns | 0.03 ns | 0.16 ns |
|  | C/N | 0.15 ns | 0.10 ns | 0.01 ns |

Values are given as F of Fisher. \*\*\*, \*\* and \* indicate significant at $p < 0.001$ , $p < 0.01$ and $p < 0.05$ , respectively. ns = not significant; I = irrigation; V = variety; D = day (D1 = 9 June, D2 = 25 June, D3 = 10 July, D4 = 1 August, D5 = 28 August); $\delta^{13}\text{C}$ = carbon isotope composition; TKN = total Kjeldahl nitrogen; C/N = carbon-to-nitrogen ratio.

Over the entire experimental period, δ^13^C values ranged from −30.67 to −28.59 ‰, with an overall increase observed at the final sampling date (+0.78 ‰ compared to earlier measurements). At D5, leaves of NG 4190 exhibited lower δ^13^C values (−29.3 ‰) than ST 4990 (−28.9 ‰) (Table 8). Total nitrogen values ranged from 0.91% to 5.28%, with a mean value of approximately 3.6% during most of the growing season, followed by a decrease at D5 (1.57%). Accordingly, the C/N ratio remained stable during the main part of the experiment (mean value of 9.56) and increased markedly at the end of the season, reaching 30.45. This pattern was driven by both increasing leaf carbon concentration and decreasing nitrogen content (data not shown).

**Table 8.**
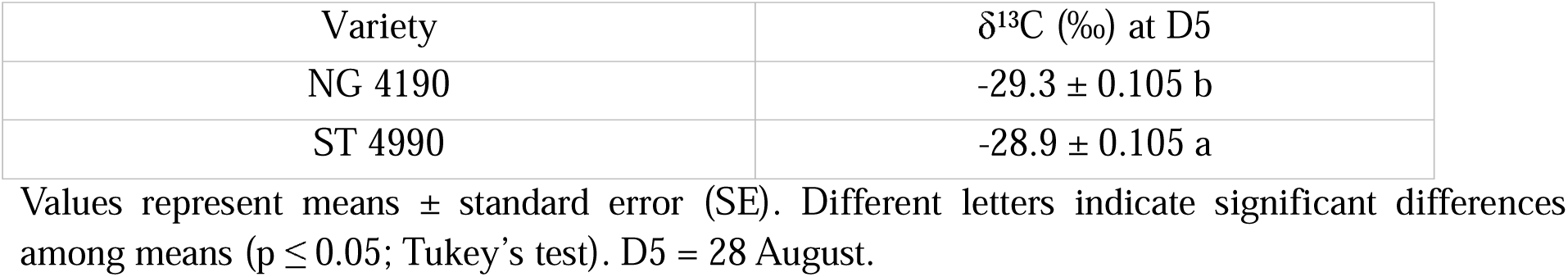
Analysis of variance (ANOVA) of interactions on carbon isotope composition (δ^13^C).

| Variety | $\delta^{13}\text{C}$ (‰) at D5 |
| --- | --- |
| NG 4190 | $-29.3 \pm 0.105$ b |
| ST 4990 | $-28.9 \pm 0.105$ a |

Values represent means ± standard error (SE). Different letters indicate significant differences among means ( $p \leq 0.05$ ; Tukey's test). D5 = 28 August.

### 3.5. Cotton yield and fiber quality indices

Table 9 presents the results of the analysis of variance (ANOVA) for yield and quality parameters. The interaction between the two factors under study (V × I) was statistically significant only for seed yield and fiber yield. In contrast, several fiber quality traits, including fiber length, uniformity index, and elongation, were not affected by the applied treatments. Micronaire (MIC) and the color parameter yellowness (+b) were strongly influenced by the irrigation regime. Specifically, MIC increased by 13% under optimal irrigation (FI), whereas the highest +b value (8.41) was observed under water deficit (DI). Fiber strength and the color parameter reflectance (Rd) were significantly influenced by variety, with the highest quality observed in ST 4990.

**Table 9.**
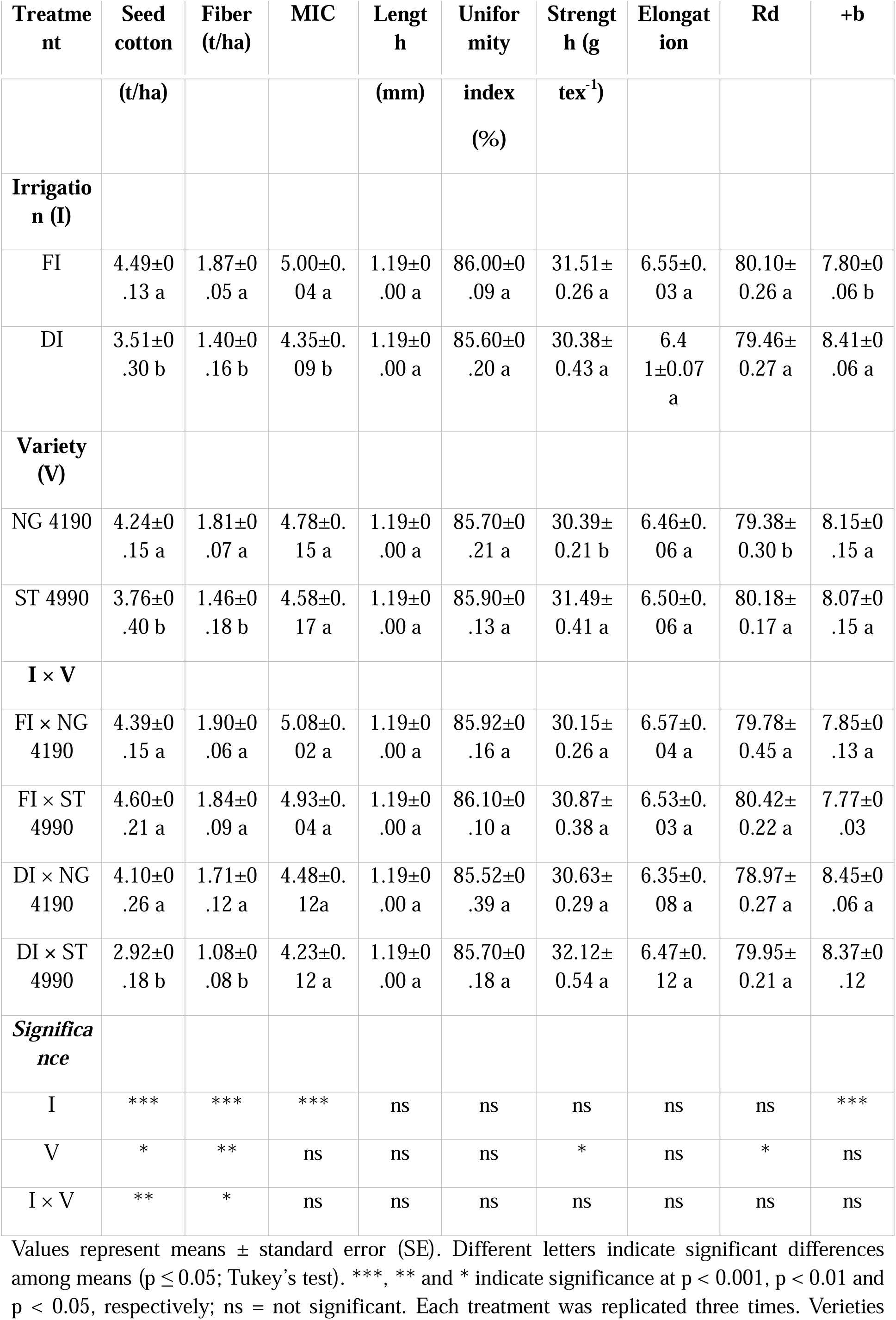
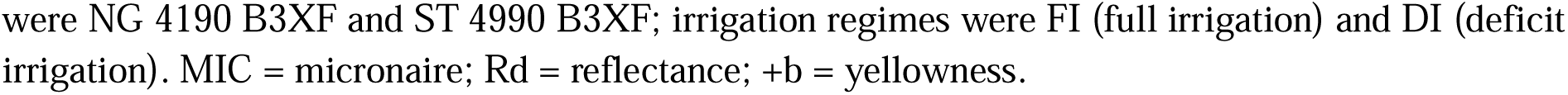
Analysis of variance (ANOVA) of main effects and interactions on cotton yield and fiber quality indices.

Regarding yield, under optimal irrigation (FI), no significant differences were observed between the two varieties, with seed yield ranging from 4.39 to 4.60 t ha^-1^ and fiber yield from 1.84 to 1.90 t ha^-1^. Under water deficit conditions (DI), NG 4190 did not show significant reductions in yield, whereas ST 4990 exhibited mean reductions of 1.68 t ha^-1^ in seed yield and 0.76 t ha^-1^ in fiber yield compared to full irrigation.

### 3.6. Correlation analysis

Pearson correlation analysis between LWP and RWC measured at predawn and midday revealed significant linear relationships (p < 0.001; Table 10). Specifically, the same parameter measured at the two times of day was strongly correlated (r ≥ 0.70). Moreover, LWP and RWC measured at the same time of day showed a strong negative correlation (r > 0.80). The only moderately strong association was observed between predawn RWC and midday LWP (r = −0.58).

**Table 10.** Pearson correlation coefficients among plant water status parameters.

|  | LWP<sub>predawn</sub> | LWP<sub>midday</sub> | RWC<sub>predawn</sub> | RWC<sub>midday</sub> |
| --- | --- | --- | --- | --- |
| LWP<sub>predawn</sub> |  |  |  |  |
| LWP<sub>midday</sub> | 0.75 |  |  |  |
| RWC<sub>predawn</sub> | -0.84 | -0.58 |  |  |
| RWC<sub>midday</sub> | -0.73 | -0.81 | 0.70 |  |

LWP = leaf water potential; RWC = relative water content. All correlations were significant at $p < 0.001$ .

In the correlation matrix shown in Fig. 9, data collected at midday (≈14:00), corresponding to the peak of water and thermal stress, revealed a high degree of consistency in the associations among physiological parameters, with statistically significant values (p < 0.001). Most correlations were strong, with the exception of a moderate association (r = 0.66) between RWC and g_s_.

**Figure 9.**
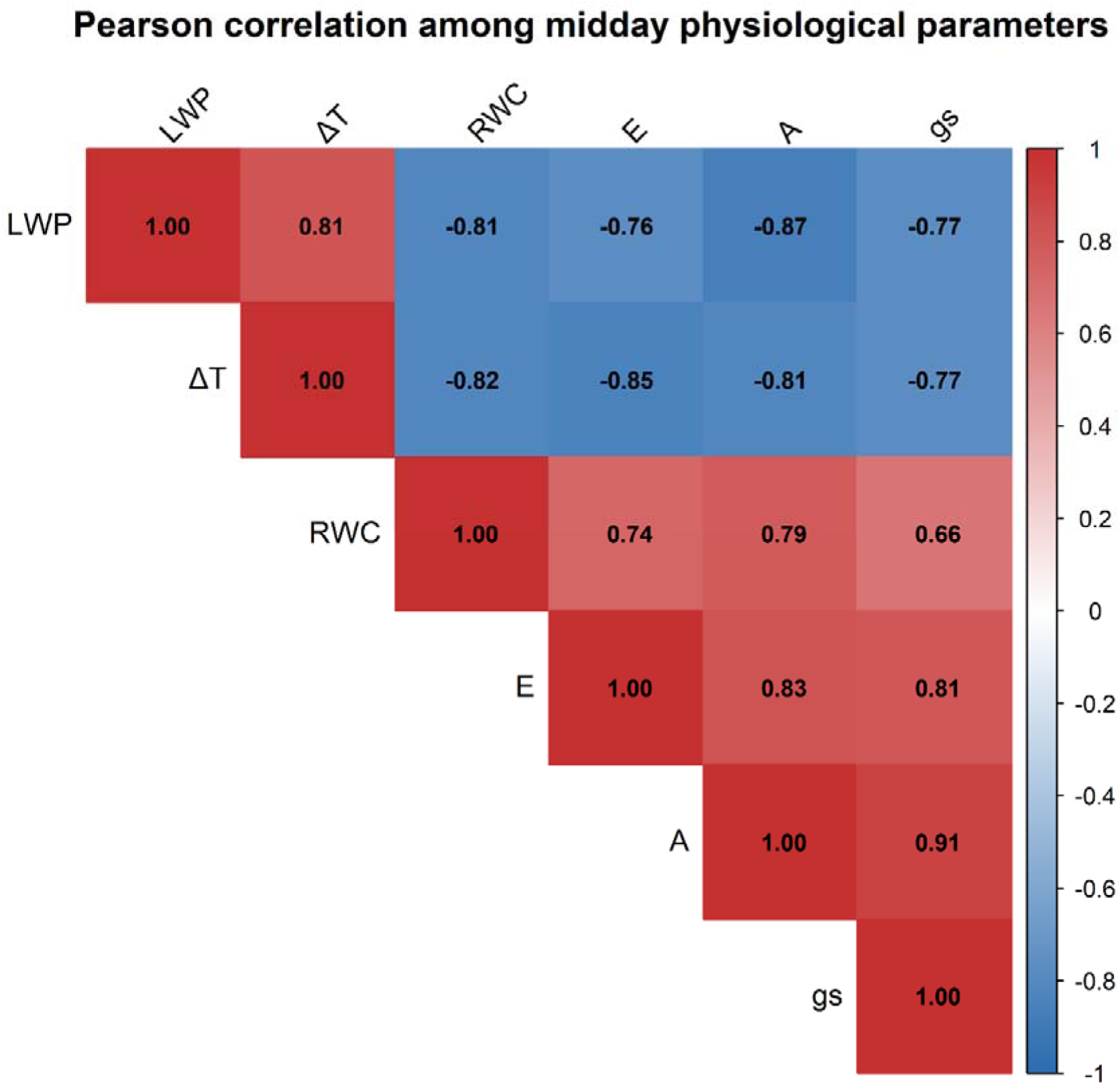
Correlation matrix of physiological parameters measured at midday. LWP = leaf water potential; ΔT = difference between canopy and air temperature; RWC = relative water content; E = transpiration rate; A = net photosynthesis rate; g_s_ = stomatal conductance. All correlations were significant at p < 0.001.

Finally, the midday (≈14:00) measurements were correlated with yield, revealing significant and strong associations, with the exception of ΔT, which showed a moderate correlation (r = −0.64) (Table 11).

**Table 11.** Pearson correlation coefficients between midday physiological parameters and total yield.

| Parameter | r (Yield) | p-value |
| --- | --- | --- |
| LWP | -0.75 | 0.005 |
| RWC | 0.76 | 0.004 |
| $\Delta T$ | -0.64 | 0.024 |
| A | 0.75 | 0.005 |
| G<sub>s</sub> | 0.73 | 0.007 |
| E | 0.76 | 0.004 |

LWP = leaf water potential; $\Delta T$ = difference between canopy and air temperature; RWC = relative water content; E = transpiration rate; A = net photosynthesis rate; g<sub>s</sub> = stomatal conductance.

## 4. Discussion

Cotton exhibits a wide genetic variability in tolerance to water deficit, which is manifested through modifications of morphological, physiological, and biochemical traits depending on the intensity of the stress. In this context, an integrated analysis of physiological responses—based on gas exchange measurements, canopy-to-air temperature differences, chlorophyll fluorescence parameters, water status indicators, and leaf chemical composition—allowed the assessment of the effects of water deficit on plant functioning and highlighted differences between the two studied varieties (NG 4190 and ST 4990) under full and moderate deficit irrigation.

The literature extensively documents the sensitivity of these indicators to water stress and their potential use for screening more tolerant genotypes (Condon et al. 2004; Baker et al. 2007; Jones 2007; de Brito et al. 2011; Guo et al. 2022; Mira-García et al. 2022). However, the results obtained in the present study indicate that, under moderate water deficit conditions, not all physiological indicators are equally effective in discriminating differences between genotypes. Specifically, only certain parameters—including midday relative water content (RWC), triose phosphate utilization (TPU), maximum photosynthesis under high irradiance (P_m_), photosynthetic efficiency at low irradiance (α), the stomatal optimization parameter g□ from the Medlyn model, and leaf carbon isotope composition (δ^13^C) measured at D5—contributed to explaining the differing productive behavior observed between the two varieties.

Under full irrigation, the two varieties did not show statistically significant differences in yield (NG 4190: 4.39 t ha^-1^ seed and 1.90 t ha^-1^ fiber; ST 4990: 4.60 t ha^-1^ seed and 1.84 t ha^-1^ fiber). However, when water supply was reduced by approximately 25%, contrasting responses were observed: NG 4190 maintained yield levels similar to those under full irrigation (4.10 t ha^-1^ seed and 1.71 t ha^-1^ fiber), whereas ST 4990 showed a marked reduction in yield, amounting to 36.5% for seed and 41.3% for fiber.

Physiological measurements taken throughout the day (gas exchange, LWP, RWC, ΔT), from predawn to midday further confirmed the importance of the timing of measurements for accurately interpreting plant responses to stress. Values recorded at dawn reflect a state of equilibrium between plant and soil water potential (Weatherley 1950), whereas measurements taken during the midday hours reflect the balance between root water supply and atmospheric evaporative demand (Williams and Araujo 2002). In the present study, treatment differences were already evident by 11:00, reaching their maximum around 14:00, when evaporative demand and transpiration generally peaked, and leaf water potential tended to reach its minimum.

The absence of significant differences in predawn measurements suggests that, despite the imposed water deficit, plants were able to reestablish water equilibrium with the soil overnight. This indicates that the applied water stress did not result in irreversible physiological damage. Consistent with this interpretation, the maximum photochemical efficiency of photosystem II (F_v_/F_m_) was higher than 0.80 in both genotypes, indicating the absence of photoinhibition and the maintenance of PSII functional integrity (Björkman and Demmig 1987; Brodribb 1996; Sunoj et al. 2025). Similar findings have been reported in other cotton studies (de Brito et al. 2011; Yi et al. 2016), where F_v_/F_m_ did not show significant changes even at leaf water potentials as low as −3.0 MPa, suggesting that this parameter may not be particularly sensitive in discriminating genotypic tolerance to water deficit. Conversely, other studies (Guo et al. 2022) have reported reductions in F_v_/F_m_ under more severe water stress, indicating that PSII response is strongly dependent on the intensity of the water deficit.

A more detailed analysis of leaf water status revealed that, with respect to values measured during the midday hours, although no significant differences in leaf water potential (LWP) were observed between the two varieties under water deficit conditions (−2.4 MPa in both), ST 4990 exhibited a slightly lower relative water content (RWC; 74.6%) compared with NG 4190 (77.7%). This result suggests that, at equivalent water potential, NG 4190 is able to maintain a higher water content in leaf tissues. Moreover, the RWC observed in NG 4190 under water deficit did not differ from that recorded in the same variety under full irrigation, despite a more negative water potential (−1.89 MPa under full irrigation).

This behavior may indicate a greater capacity to maintain cellular turgor, potentially associated with osmotic adjustment and/or differences in the elastic properties of leaf tissues (Jones 1990; Clifford et al. 1998; Li et al. 2022). Consistently, the absence of significant differences in stomatal conductance between varieties during the midday hours suggests that the observed differences in tissue water content are not directly attributable to variations in instantaneous stomatal regulation. Under full irrigation, ST 4990 displayed slightly less negative LWP values (on average ∼0.34 MPa) compared with NG 4190, despite similar RWC and gas exchange between the two varieties.

Although more negative water potentials may reflect differences in soil–plant hydraulic conductance (Jones 1990; Steudle and Peterson 1998), the similarity in RWC and gas exchange supports the hypothesis that NG 4190 has a greater ability to maintain tissue water content even at lower water potentials.

The absence of significant differences in gas exchange measurements (A, E, g_s_) suggests that such instantaneous assessments may not fully capture the physiological processes underlying yield differences, particularly under moderate water stress. These measurements represent a snapshot of leaf physiological status and do not necessarily reflect integrated plant responses over time (Medrano et al., 2015; Leakey et al. 2019).

Supporting this observation, correlation analysis between carbon isotope composition (δ^13^C), the internal-to-ambient CO□ ratio (C_i_/C_a_), and instantaneous water-use efficiency (iWUE, A/g_s_) did not reveal strong associations (data not shown), contrasting with reports in the literature (Saranga et al. 1998; Condon et al. 2004; Du et al. 2021). Similarly, canopy-to-air temperature differences (ΔT) did not show significant variation between varieties, suggesting similar instantaneous transpiration behavior.

Statistical differences in instantaneous gas exchange were observed only at the second measurement date (25 June; data not shown) for stomatal conductance (g_s_). Under water deficit, NG 4190 exhibited higher values than ST 4990, likely reflecting increased stress severity at this sampling date. This confirms that the sensitivity of instantaneous measurements strongly depends on stress intensity. Previous studies have shown that net photosynthesis declines rapidly only when water deficit becomes more severe, concomitant with a marked reduction in stomatal conductance below approximately 0.4 mol m^-2^ s^-1^ (Baker et al. 2007).

In contrast to instantaneous measurements, carbon isotope composition (δ^13^C) provides an integrated indicator of whole-plant water-use efficiency over time, reflecting long-term stomatal behavior and carbon assimilation dynamics (Condon et al. 2004). In the present study, differences in δ^13^C were observed only at the end of the growing cycle, near harvest, without showing a clear response to irrigation treatment, and were primarily associated with genotypic variability. Specifically, NG 4190 exhibited more negative δ^13^C values than ST 4990, reflecting a greater discrimination against the heavier carbon isotope and a greater stomatal openness (see Fig. 5J) to fascilitate CO_2_ diffusion to the carboxylation site (de Brito et al. 2011; Sushma et al. 2024).

The emergence of such differences in the final stages of the cycle reflects the dynamic nature of δ^13^C, which can vary over time, particularly during boll maturation, in response to the cumulative effect of late-season water stress on intrinsic water-use efficiency (iWUE) (Saranga et al. 1999; Leakey et al. 2019). In contrast, total nitrogen content (TKN) and C/N ratio followed the typical seasonal trend, without significant differences between irrigation treatments. This indicates that moderate water deficit did not alter the normal dynamics of leaf nitrogen remobilization.

Furthermore, no significant differences were observed in the main biochemical parameters derived from A-Ci curves, such as the maximum Rubisco carboxylation rate (V_cmax_), the maximum electron transport rate (J_max_), and dark respiration (R_d_). In contrast, genotypic differences emerged in parameters derived from light response curves, particularly maximum photosynthesis under high irradiance (P_m_) and photosynthetic efficiency at low irradiance (α). Consistent results were also obtained from chlorophyll fluorescence rapid light curves (RLCs): under water deficit, NG 4190 exhibited slightly higher initial PSII photochemical efficiency (α_PSII_) compared with ST 4990, whereas the maximum electron transport rate (ETR_max_) was influenced solely by irrigation level and not by genotype.

Overall, these results indicate that the observed yield differences cannot be attributed to variations in the intrinsic biochemical capacity of the photosynthetic apparatus in individual days, but rather to the differences in the longer term physiological regulation of photosynthetic performance. In line with this interpretation, factors such as plant water status (midday RWC) and optimal stomatal behavior described by the g□ parameter of the Medlyn model may have played an important role in modulating photosynthetic performance under moderate water deficit conditions.

This interpretation is supported by previous studies indicating that stomatal behavior is not directly regulated by Rubisco activity, as reductions in carboxylation capacity do not necessarily lead to changes in stomatal conductance (von Caemmerer et al. 2004; Medlyn et al. 2011). Moreover, it has been reported that, under moderate water stress, stomatal limitations tend to dominate over non-stomatal limitations, whereas as leaf water potential decreases (below approximately −1.5 MPa), non-stomatal limitations progressively become more relevant (Faver et al. 1996). However, the results of the present study demonstrate that, even at water potentials around −2.4 MPa, non-stomatal limitations did not constitute the main controlling factor of photosynthetic processes in the two analyzed varieties.

An additional distinguishing factor between the genotypes is the stomatal optimization parameter g□ from the Medlyn stomatal model, which is directly related to the marginal water cost associated with carbon assimilation (Medlyn et al. 2011). Higher g□ values indicate a less conservative stomatal strategy, characterized by greater water loss per mole of CO□ assimilated. In the present study, under water deficit conditions, NG 4190 exhibited higher g□ values than ST 4990 (5.99 vs. 5.30), suggesting a greater propensity for leaf gas exchange.

Although higher stomatal conductance is generally associated with reduced water-use efficiency (WUE), under moderate water deficit this behavior can be advantageous in terms of cumulative carbon assimilation. Greater stomatal opening can allow leaves to more effectively exploit temporal variations in canopy irradiance, such as sunflecks, which occur on timescales ranging from seconds to several hours (McAusland et al. 2016). Because stomatal closure limits CO□ availability at the carboxylation site and reduces photosynthetic assimilation (Chaves 1991), a less conservative stomatal strategy can thus sustain higher carbon assimilation over time, with potential positive effects on productivity (Medrano et al. 2015; Luo et al. 2016).

The ability of NG 4190 to maintain similar yield levels under full irrigation and water deficit may also be supported by differences in resource allocation between vegetative and reproductive growth. Water-use efficiency (WUE), expressed as the ratio between harvestable yield and water consumed, can vary even in the absence of changes in total carbon assimilation or overall water use if modifications occur in the harvest index (HI) (Leakey et al. 2019; Conaty and Constable 2020).

Correlation analysis between LWP and RWC measured at predawn and midday revealed strong linear relationships (r ≥ 0.70; p < 0.001), indicating that both times of day reflect similar trends in water status parameters. However, predawn measurements did not capture genotypic differences, suggesting that for studies aimed at distinguishing genotypic responses, midday measurements are more informative and sufficient, while also reducing experimental workload. From the analysis of physiological parameters measured at midday, the only moderate association observed between RWC and g_s_ confirms that leaf water content is not determined solely by instantaneous stomatal regulation, but also reflects other physiological mechanisms and plant characteristics such as tissue hydraulic conductivity. Overall, physiological parameters measured at midday closely correlate with yield, indicating that they effectively represent crop performance.

Finally, the analysis of fiber technological traits indicated that, in general, moderate water deficit does not significantly alter fiber quality, as many parameters are primarily under genetic control, except under more severe stress conditions (de Brito et al. 2011; Chalise et al. 2022). Some exceptions were observed: micronaire (MIC) tended to decrease under water deficit, since higher irrigation promotes better cellulose deposition from seeds to fiber (Manibharathi et al. 2024), while the yellowness index (+b) was slightly increased. In our study, ST 4990 exhibited slightly higher fiber strength and Rd color values compared with NG 4190, traits reflecting, respectively, the mechanical robustness of the fiber and its lighter hue, both important for industrial processing and final textile quality. Both genotypes, however, possess high-quality fibers, with strength exceeding 30 g tex^-1^ and Rd around 80, values consistent with high commercial standards (Bradow and Davidonis 2000).

Some limitations should be considered when interpreting these results. The study did not include direct measurements of root architecture, resource allocation, or osmolyte levels, factors that could influence responses to water deficit. Furthermore, the investigation was conducted over a single growing season, although the experimental design with adequate replication provides a solid statistical basis. Finally, the factors underlying variability in the g□ parameter warrant further investigation; it would be useful to explore potential correlations with leaf traits or other physiological characteristics, and to identify possible markers associated with this parameter, as understanding of the mechanisms driving g□ variability remains incomplete in the literature.

## 5. Conclusion

The results of this study conducted in Texas allowed the identification of physiological markers capable of discriminating cotton genotypes tolerant to moderate water stress. These findings suggest that differences in photosynthetic performance primarily emerge under dynamic, fluctuating light conditions rather than under steady-state conditions. Furthermore, the applied moderate water deficit did not compromise photosystem integrity, confirming the absence of irreversible damage in the plants.

Analysis of carbon isotope composition (δ^13^C) confirmed time-integrated genotypic differences in water-use efficiency, with NG 4190 characterized by more negative values, consistent with a less conservative stomatal strategy and greater stomatal opening. Between the two studied varieties, NG 4190 exhibited greater yield stability under moderate water deficit, maintaining seed and fiber production levels comparable to those observed under full irrigation. In contrast, ST 4990 showed a reduction in yield but displayed slightly superior fiber technological properties, with stronger fibers and a lighter color (Rd).

The data support our hypothesis that differences in yield stability under moderate water stress are primarily associated with variation in leaf photosynthetic traits and stomatal behavior.

Overall, our data suggest that RWC_midday_, P_m_, α, g□, and δ^13^C emerge as the most promising physiological targets for breeding programs aimed at improving drought tolerance in cotton, highlighting that selection of the appropriate genotype can save water without significant yield loss or quality compromise. However, further validation across multiple seasons and a wider range of genotypes is needed to confirm the robustness and general applicability of these findings.

## Author Contributions

N.T. conducted experiment, curated and analyzed data, wrote and revised the manuscript. X.D. conceived idea, designed, supervised and conducted experiment, curated and analyzed data, and revised the manuscript. T.F.D. designed and conducted experiment, curated data, and reviewed and revised the manuscript. N.I. supervised data analysis, reviewed and edited the manuscript. U.A. conducted experiment, reviewed and edited the manuscript. T.T. valided data analysis, reviewed and edited the manuscript. All authors approved the final form of the manuscript.

## Acknowledgments

We appreciate Randy Cox, Gabe Diaz, Manuel Figueroa Pagan and Joe Gonzalez for assistance in field equipment instellation and crop management. We also appreciate Christine Thompson and Liza Silva for administrative support. We thank Dale Mott for providing cotton seed and allowing us to use his gin machine, David Kerns for offering advice on thrip control, and Charlotte Daniels for conducting leaf nitrogen analysis.

## Funding

X.D. received funding from the Cotton Incorporated (project 20-557TX) and USDA NIFA (Hatch project TEX0-1-9574). D.F.T. received funding from Brazilian Federal Agency for Support and Evaluation of Graduate Education (CAPES/ PRAPG, Notice 14/2023). N.T. received funding from University of Palermo, Italy to visit Uvalde Research Center.

## Conflict of Interest Statement

The authors declare no conflicts of interest.

## Data Availability

Data files and further details of merthods supporting this study are available as Supplementary material at https://zenodo.org/records/19864799.

## Notes

https://zenodo.org/records/19864799

### Competing Interest Statement

The authors have declared no competing interest.

